# *Laccaria bicolor* pectin methylesterases are involved in ectomycorrhiza development with Populus tremula x Populus tremuloides

**DOI:** 10.1101/2022.06.08.495362

**Authors:** Jamil Chowdhury, Minna Kemppainen, Nicolas Delhomme, Iryna Shutava, Jingjing Zhou, Junko Takahashi, Alejandro G. Pardo, Judith Lundberg-Felten

## Abstract

- The development of ectomycorrhizal (ECM) symbioses between soil fungi and tree roots requires modification of root cell-walls. The pectin-mediated adhesion between adjacent root cells loosens to accommodate fungal hyphae in the Hartig Net, facilitating nutrient exchange between partners. We investigated the role of fungal pectin modifying enzymes in *Laccaria bicolor* for ECM formation with *Populus tremula* x *Populus tremuloides*.
- We combine transcriptomics of cell-wall related enzymes in both partners during ECM formation, immunolocalization of pectin (Homogalacturonan, HG) epitopes in different methylesterification states, pectin methylesterase (PME) activity assays and functional analyses of transgenic *L. bicolor* to uncover pectin modification mechanisms and the requirement of fungal pectin methylesterases (*LbPMEs*) for ECM formation.
- Immunolocalization identified remodelling of pectin towards de-esterified HG during ECM formation, which was accompanied by increased *LbPME1* expression and PME activity. Overexpression or RNAi of the ECM-induced *LbPME1* in transgenic *L. bicolor* lines led to reduced ECM formation. Hartig Nets formed with *LbPME1* RNAi lines were shallower, whereas those formed with *LbPME1* over-expressors were deeper.
- This suggests that *LbPME1* plays a role in ECM formation potentially through HG-de-esterification, which initiates loosening of adjacent root cells to facilitate Hartig Net formation.

## Introduction

Ectomycorrhizal (ECM) symbiosis is one of the most predominant forms of plant-microbe interactions in boreal and temperate forest ecosystems (Read *et al*., 2004; Qu *et al*., 2010; McGuire *et al*., 2013) where it plays a key role in plant health through mineral nutrient cycling. During symbiosis establishment, ECM fungi form a sheath around the lateral root tips and colonize the apoplastic space between epidermal and cortical root cells (Kottke & Oberwinkler, 1987; Abras *et al*., 1988), forming the Hartig Net, a nutritional exchange site between plant and fungus. Formation of this symbiotic interface requires modification of plant and fungal cell-walls (Brundrett, 2004; Martin *et al*., 2017) and has interested researchers for more than 30 years (Paris *et al*., 1993; Balestrini *et al*., 1996a; Balestrini & Bonfante, 2014) . I*n situ* immuno-labelling-based characterization of the cell-wall polymer epitopes (pectin, cellulose, hemicellulose) in mature ECM in comparison to non-mycorrhizal roots, has underpinned our understanding of how the plant cell-wall is altered during the establishment of hyphae in the apoplastic space. The suggested key mechanisms for Hartig Net formation are (i) mechanical force from the hyphal tip (Garcia *et al*., 2015), (ii) auxin secretion from root tips to promote cell-wall loosening (Gea *et al*., 1994) and (iii) secretion of fungal cell-wall-degrading enzymes (Bonfante & Genre, 2010; Veneault-Fourrey *et al*., 2014). The two former hypotheses await further proof through functional studies, whereas the latter has largely benefitted from the availability of fungal genomes and the possibility of using RNAi techniques in the ectomycorrhizal model fungus *L. bicolor* to identify and functionally characterize potential fungal plant cell-wall degrading (PCWDE) enzymes (Zhang *et al*., 2018, 2021) . Whole-genome sequences of several ECM species have revealed that ECM fungi have a reduced set of PCWDEs compared to their saprotrophic ancestors (Martin *et al*., 2008; Kohler *et al*., 2015; Miyauchi *et al*., 2020). ECM fungi likely employ these PCWDEs to modify the plant cell-wall matrix for apoplastic accommodation so to establish bi-directional nutrient transport. A significant increase of *PCWDE* expression during ECM interactions further supports this hypothesis (Sebastiana *et al*., 2014; Veneault-Fourrey *et al*., 2014; Kohler *et al*., 2015).

The extracellular route that fungal hyphae take during Hartig Net formation is rich in pectic polysaccharides; in particular, homogalacturonan (HG) is predominant in the middle lamella between adjacent root cells (Daher & Braybrook, 2015). Pectic polysaccharides make up to 35% of the total cell-wall dry weight and play versatile roles in plant physiological processes including cell growth and differentiation. Their dynamic alterations can modify cell-wall chemistry and rheology (Bidhendi & Geitmann, 2016). HG modifying enzymes (HGMEs) play a central role in cell-to-cell adhesion and cell separation (Sénéchal *et al*., 2014; Daher & Braybrook, 2015) and include multigenic families such as pectin methylesterases (PMEs), pectin acetylesterases (PAEs), pectate lyases (PLs), and polygalacturonases (PGs) (Sénéchal *et al*., 2014). The ECM basidiomycete *Laccaria bicolor* has a restricted set of HGMEs comprising only four PMEs and six PGs (Martin *et al*., 2008), of which the ECM-induced *LbGH28A* has recently been functionally characterized for its activity on pectin and polygalacturonic acid and is suggested to contribute to Hartig Net formation (Zhang *et al*., 2022). On the other hand, its host plant *Populus tremula* has a large set of HGMEs comprising 73 *PMEs*, 12 *PAEs*, 28 *PLs* and 38 *PGs* (Sundell *et al*., 2015). Despite the reduced *HGME* set, *L. bicolor HGMEs*, i.e. *LbPME*s and *LbPGs*, are transcriptionally up-regulated at various time-points during an interaction with *Populus* ((Veneault-Fourrey *et al*., 2014; Kohler *et al*., 2015) and the current study) suggesting an involvement of these *HGMEs* in ECM development. PMEs seem to act as first modifiers of HG modification (Pelloux *et al*., 2007; Manmohit Kalia, 2015; Sénéchal *et al*., 2015) and catalyse the de-esterification of the C6-linked methyl-ester groups of HG chains. Although the exact modes of action of PMEs are still debated, PME-mediated HG de-esterification patterns can be random or block-wise, with enzyme isoforms and cell-wall pH being determining factors (Catoire *et al*., 1998; Denès *et al*., 2000; Kim *et al*., 2005). Thus, depending on physiological conditions, PME activity regulates both cell-wall loosening and cell-wall stiffening (Micheli, 2001; Sénéchal *et al*., 2014). During cell-wall loosening, PME de-esterification activity makes HG more accessible to depolymerizing enzymes like PLs and PGs (Micheli, 2001; Pelloux *et al*., 2007; Sénéchal *et al*., 2014; Manmohit Kalia, 2015). As it has recently been shown that a fungal PG (LbGH28A) is involved in Hartig Net formation (Zhang *et al*., 2022), we are here hypothesizing that preceding steps of HG modification also receive a fungal contribution. Several fungal pathogens employ PMEs to overcome pectin-rich cell-wall barriers (Lionetti *et al*., 2012; Sella *et al*., 2016; Fan *et al*., 2017). ECM fungi encoding PMEs may apply similar strategies to modify cell-walls to form the Hartig Net. We hypothesize that *L. bicolor* PMEs facilitate cell-wall loosening during ECM development. Therefore, we investigated the functional role of *L. bicolor* PMEs in ECM symbiosis. Using *L. bicolor* and *P. tremula* x *P. tremuloides* interactions in an *in vitro* culture system, we leveraged whole-genome transcriptomics, microscopy coupled with immunolabeling and transgenic approaches targeting *LbPMEs* to explore the methylesterification state of plant cell-walls in ECM and the potential role of *L. bicolor PMEs* during ECM formation.

## Materials and methods

### Fungal strains and growth conditions

*L. bicolor* (Maire) isolate S238N, empty vector lines EV7 and EV9 (Plett *et al*., 2011) and transgenic lines (this study) were maintained on agar Pachlewski P5 medium at 20°C. For experiments, we used 14-days old free-living mycelia (FLM) grown on cellophane placed on solidified Pachlewski P20 medium. For media and culture conditions see (Felten *et al*., 2009). *Populus tremula* x *Populus tremuloides* (T89) cuttings were micro-propagated under *in vitro* conditions and grown on half strength MS medium (Murashige & Skoog, 1962) containing 2% sucrose and 1% plant agar (Cat # P1001, Duchefa Biochemie B.V, The Netherlands) with pH 5.8 in rectangular 14 x 7.5cm, 7.5 cm high pots under a 16 h photoperiod at 22° C. After one month, rooted cuttings were transferred to 12cm square Petri dishes containing freshly prepared half strength MS medium. The root systems were covered above and below with a cellophane membrane and grown vertically for two weeks before we started interaction studies through an *in vitro* sandwich culture system on MES-buffered P20 medium with cellophane-grown FLM as described in Felten *et al*. (2009).

### RNA extraction and cDNA synthesis

The time-course samples for RNA-sequencing data were derived from the *in vitro* sandwich culture of *L. bicolor* interacting with *P. tremula* x *P. tremuloides* as mentioned above. Control plant and fungal samples were grown in separate plates without any interactions. At 3, 7, 14, 21 and 28 days after fungal contact (DAC) ten root tips (ECM or control) each about 0.5 cm long, were harvested from two plants per condition and pooled to be considered as one replicate. We analysed four such replicates per time-point and condition. For free-living mycelia controls, young hyphae were collected at the periphery of fungal cultures without plants. Total RNA was extracted with RNAqueous™ Total RNA Isolation Kit (Invitrogen) following the manufacturer’s instructions, except that 1% polyvinylpyrrolidone (PVP) was added to the extraction buffer. The remaining gDNA was removed by DNA-free™ DNA Removal Kit (Cat. # AM1906, ThermoScientific, USA) followed by purification of the RNA by RNeasy MinElute Cleanup Kit (Cat # 74204, Qiagen). RNA concentration and quality were checked by Nanodrop 2000 spectrophotometer (Thermo Scientific) and by Bioanalyzer RNA 6000 pico chip (Cat. # 5067-1513 Agilent.com, USA), respectively. The full-length cDNAs were generated according to the Smart-seq2 protocol (Picelli *et al*., 2014) followed by quality control analysis with Agilent High Sensitivity DNA Kit (Cat. # 5067-4626, Agilent.com, USA). Sequencing libraries were generated using a Rubicon ThruPLEX DNA-seq kit (Takara Bio USA, Inc.) and sequencing of all 55 samples took place on a single Illumina NovaSeq6000 S2 flow-cell lane generating 150bp paired-end reads at SciLifeLab (Science for Life Laboratory, Stockholm, Sweden).

For qPCR-analysis the procedures for RNA extraction from free-living mycelia, cDNA synthesis, conditions for qRT-PCR and gene expression analysis are identical to those described in Kemppainen et al. (2020). Four replicates per condition and genotype were assessed by qPCR analysis, except for the initial large-scale screening of transgenic fungal lines, where only one biological replicate per line was analyzed. The primer sequences used in the qRT-PCR expression analysis are provided in Table **S1**.

### Pre-processing of RNA-Seq data and differential expression analyses

Data pre-processing was performed as described by (Delhomme *et al*., 2014). Details can be found in Method **S1**. The raw data are available from the European Nucleotide Archive (ENA, https://ebi.ac.uk/ena) under the accession number PRJEB41173.

### Pectin methylesterase activity assay

PME activity in the ECM and control plant root tips was measured in a coupled enzyme-based spectrophotometric assay (Grsic-Rausch & Rausch, 2004) with a spectrophotometric 96-well plate reader (Epoch, Biotek, USA). Starting material comprised 50 mg (FW) from each of five biological replicates of root/mycorrhizal samples. We measured PME activity in the FLM, following the method described by Anthon et al. (2004) with modifications to increase sensitivity and consistency (Method **S2**).

### Sample preparation for microscopy and immunolocalization

The ECM and control root tips of about 0.5 cm length were collected at 7, 14, 21 and 28 days after fungal contact (DAC). At each time-point, we collected 10 to 12 root tips from five to six plants, fixed them in 4% paraformaldehyde in PBS overnight at 4 °C, infiltrated and embedded them in LR white resin (hard grade) and used a glass knife microtome to make 1µm thick cross-sections following the protocol described by Burton et al. (2011). For immunolabeling, we localized pectin epitopes with antibodies (LM19 and LM20, that preferentially recognize de-esterified or highly methylesterified HG epitopes, respectively (Verhertbruggen *et al*., 2009)) on three sequential cross-sections (considered as equivalent sections) cut at around 500 µm from the root apex following the protocol described by Chowdhury et al. (2014) except that we used the Cy™5 conjugated anti-rat secondary antibody and Alexa Fluor 488 conjugated with WGA for FLM labeling (Method **S3**). For negative controls, sections were treated similarly except that the primary antibodies were not added. No fluorescence was visible (images were black) in negative controls. For ECM anatomy analysis, we used double stainings with Alexa Fluor 488 conjugated with WGA (10µg/ml final concentration) to reveal fungal tissues and, for the plant cell-wall, 0.01% Pontamine fast scarlet 4BS in PBS (Anderson *et al*., 2010). For transmission electron microscopy, we localized pectin epitopes with LM19 and LM20 antibodies in 70 nm thick resin embedded sections, which were recognized by 5nm gold conjugated anti-rat IgG secondary antibody (Wilson & Bacic, 2012) and visualized with a Talos L120C Transmission Electron Microscope. Gold beads were quantified manually on 10-16 images from three individual root sections per antibody and condition.

### Confocal image acquisition and image analysis

Images were acquired with a Confocal Laser Scanning Microscope (Zeiss LSM780) equipped with a x40 1.2 NA water immersion objective. The Alexa Fluor 488 conjugated with WGA and the Cy™5 conjugate with secondary antibody were excited with 488 nm and 633 nm laser lines and detected at 500 to 590 nm and 640 to 750 nm, respectively. We indirectly quantified pectin epitopes by measuring the fluorescence probe (Cy5) intensity (mean number of pixels per unit area) conjugated with the respective primary antibody using the ‘curve spline’ function of the Zeiss ZEN blue software. The intensity values were normalized by subtracting the background obtained from negative controls of equivalent sections.

### PME vector construction and generation of *L. bicolor* PME transgenic lines

We created RNAi constructs driven by the constitutive *Agaricus bisporus gpdII* promoter and silencing either *LbPME1* (*LbPME1*_RNAi) or simultaneous silencing of *LbPME1* and three highly similar *Laccaria* PMEs: *LbPME2* (JGIv2 ID #676331), *LbPME3* (JGIv2 ID #315258), *LbPME4* (JGIv2 ID #315313) (referred to as *LbPME1-4*_double RNAi) (Fig. **S1**, **S2**). *L. bicolor* cloning and transformation (Kemppainen *et al*., 2005, Kemppainen & Pardo, 2010), selection and insert number analysis by ddPCR (Głowacka *et al.,* 2016) and insertion site characterization (Table **S2**) using plasmid rescue (Kemppainen *et al.,* 2008) and TAIL PCR (Liu *et al.,* 1995, Liu *et al.,* 2007) are described in detail in Methods **S4-S7**.

## Results

### Time-course gene expression profiling reveals alteration of cell-wall modifying genes during *L. bicolor and P. tremula* x *P. tremuloides* interactions

We performed a time-course gene expression study by RNA-Seq to reveal differentially expressed genes (DEGs) related to plant cell-wall modification in *P. tremula x P. tremuloides* and *L. bicolor* at 3, 7, 14, 21 and 28 DAC in colonized lateral root tips or the respective controls (control root tips without fungus or free-living mycelium) (Tables S3, S4). These time-points during ECM formation correspond to hyphal adhesion on the root (0-3 days), mantle formation (3-7 days), penetration of the root (7-14 days) and Hartig Net maturation (21-28 days) (Fig. **S3**), as revealed by microscopic observation. *L. bicolor* and *P. tremula* x *P*. *tremuloides* cell-wall-related transcriptome profiles were markedly influenced by their interaction (Figs **1**, **S4**, **S5**). A higher proportion of plant cell-wall-degrading enzymes (PCWDEs) encoding genes related to pectin, hemicellulose and cellulose, were up-regulated in the fungal genome (up to 68% up-regulated genes in the respective family) as compared to the plant genome (up to 12% up-regulated genes in the respective families) (Figs **2**, **S5**). None of the xylan modifying enzymes present in *L. bicolor* or *Populus* was induced. Several Homogalacturonan (HG) modifying enzymes were differentially expressed in both plant and fungus (Figs **3**, **S6**). This included the specific up-regulation of three out of 73 *Populus PMEs* at 3, 7 and 14 DAC and down-regulation of seven *Populus PMEs* at 21 and 28 DAC. Only one of the four *L. bicolor PMEs* (*LbPME1)* showed significant up-regulation at all time-points (confirmed by qPCR, Fig. **S7**) while none of the others showed any changes (Figs **3**, **S6**, **S8**). Our data also revealed differential expression of *Populus PME inhibitors (PMEIs),* an inhibitory protein family of plant PMEs (Di Matteo *et al*., 2005; Wormit & Usadel, 2018), only during the late stages, at 21 DAC and 28 DAC, with two members being up-regulated, and three members down-regulated (Fig. **S6**). Plant PMEIs are unlikely to inhibit fungal PMEs due to the lack of conserved critical residues in fungal PMEs for a PME-PMEI interaction (Giovane *et al*., 2004; Di Matteo *et al*., 2005; Lionetti *et al*., 2007; Reca *et al*., 2012). Protein alignments confirmed the absence of the PMEI domain (PF04043) in *L. bicolor* PMEs (and a wide range of fungi with different lifestyles) (Fig. **4**), suggesting that post-translational inhibition of fungal PMEs by plant PMEIs is an unlikely event. The PME domain structure occurs as a contributing factor for phylogenetic clustering when plant and fungal PMEs are aligned (Fig. **4a**).

**Fig. 1:**
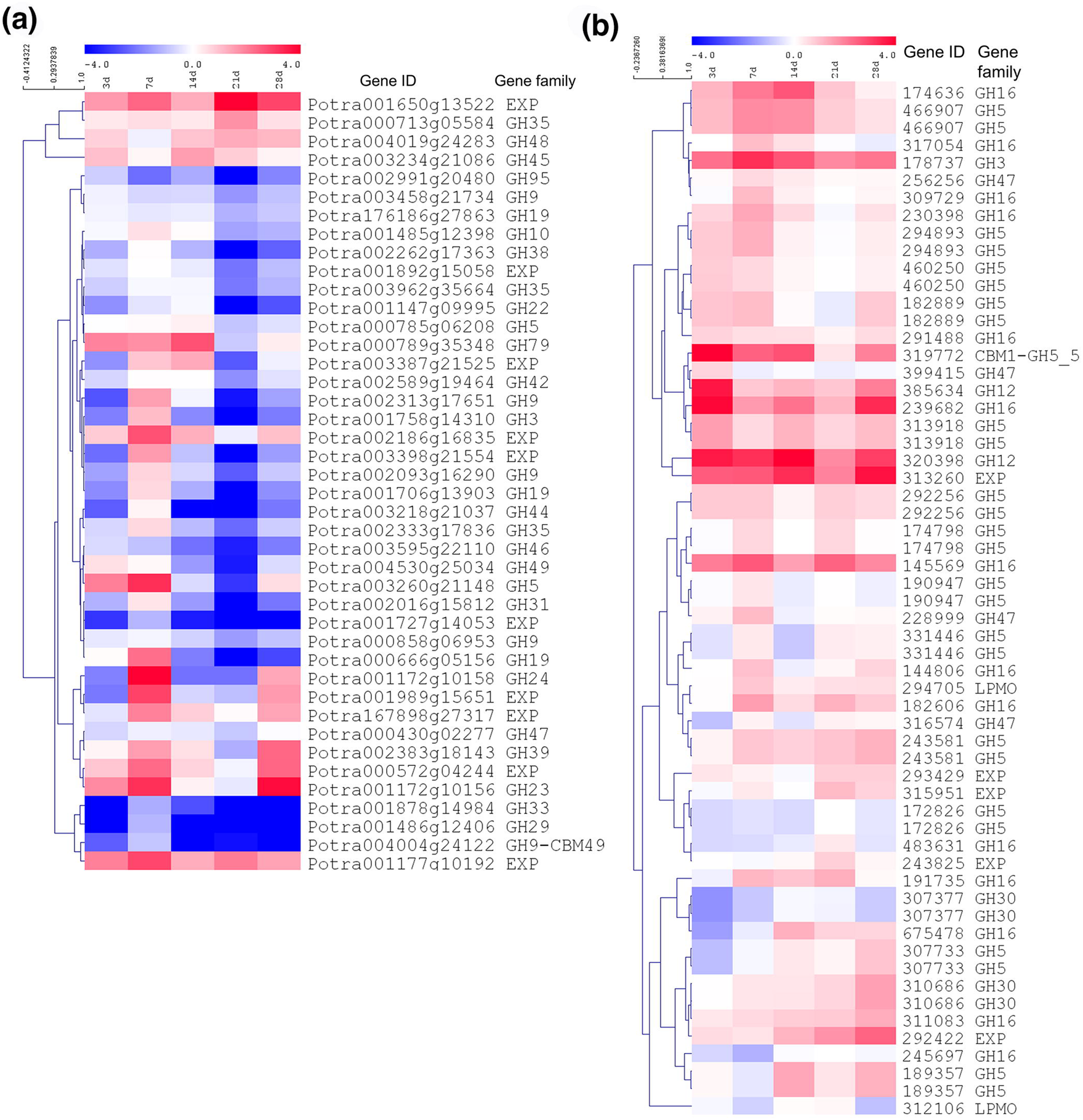
Heat map of relative gene expression levels of Differentially Expressed Genes (DEGs) related to cellulose and hemicellulose degrading enzymes in (a) *P. tremula* x *P. tremuloides* and (b) and *L. bicolor*. Expression matrix of log_2_ fold change values cluster for gene based on Pearson’s clustering method.

**Fig. 2:**
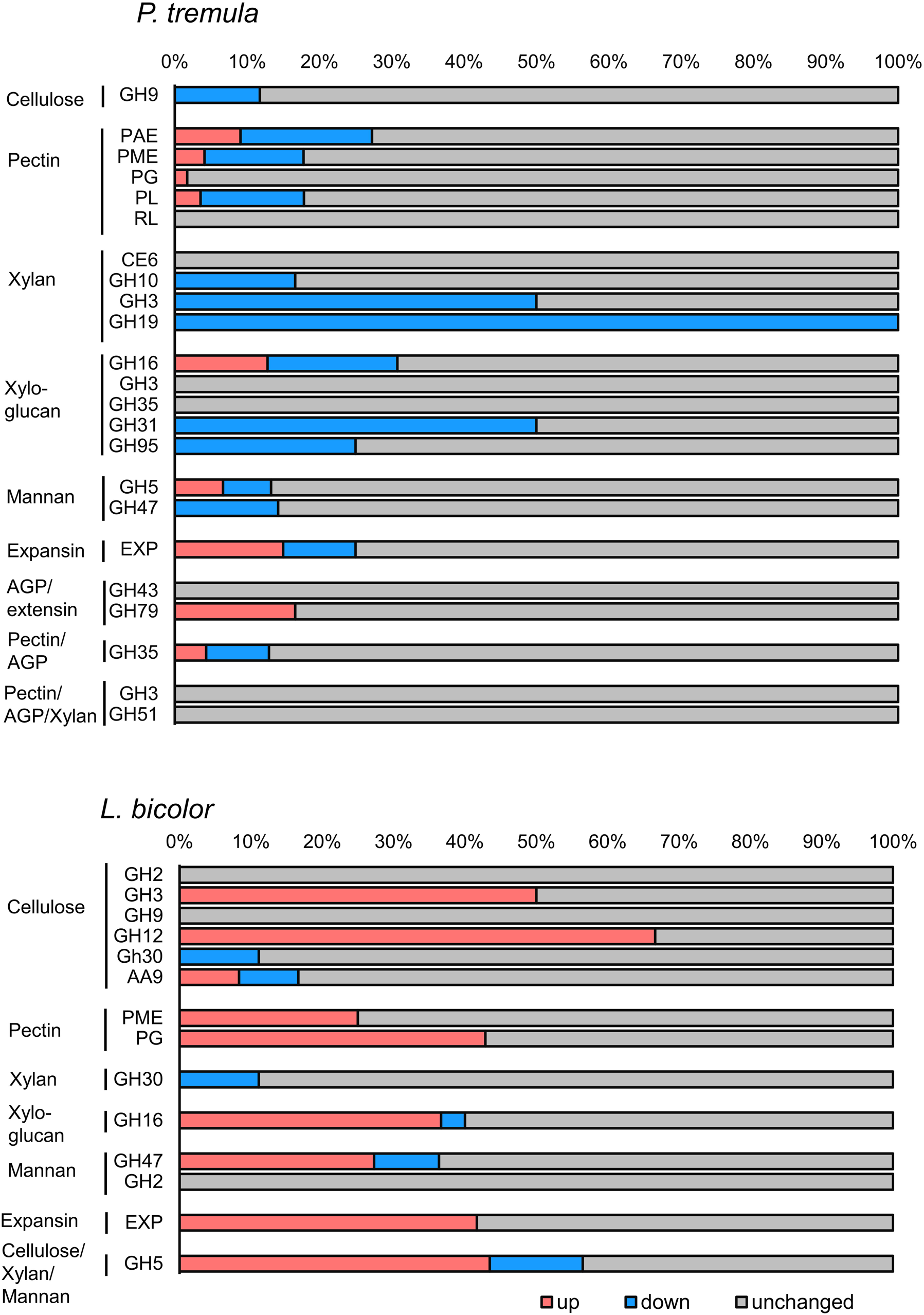
Proportion of differentially expressed and unchanged cell-wall-degrading enzymes (CWDEs) among key gene families in *P. tremula* x *P. tremuloides* and *L. bicolor* potentially play crucial roles during ECM formation. Plant genes were selected based on *Populus* whole-genome annotation (Sundell *et al*., 2015) and functional analyses based on RNA expression data (Kumar *et al*., 2019). Fungal genes are annotated from JGI database.

**Fig. 3:**
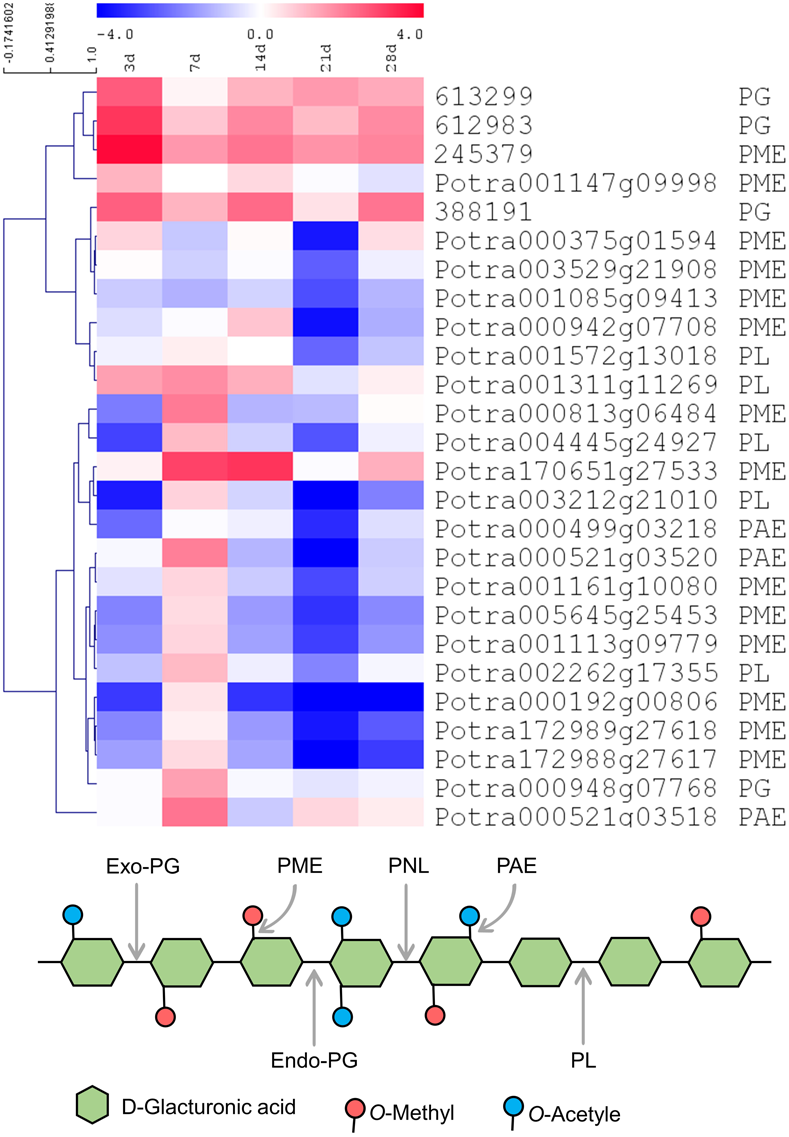
Time course RNA-Seq expression profile of the potential candidate homogalacturonan (HG) modifying enzymes (HGMEs) during *L. bicolor* interaction with *P. tremula x P. tremuloides*. The heat map represents log_2_ fold change ratios of differentially regulated genes (log_2_ fold change 0.5, padj<0.05) in colonized roots versus control roots without fungus harvested at the respective similar time-point. Fold change values are indicated. PME = Pectin methylesterase, PMEI = Pectin methylesterase inhibitor, PAE = Pectin acetylesterase, PL = Pectate lyase, PG = Polygalacturonase, PGI = Polygalacturonase inhibitor. Note that *PMEI, PAE, PGI*, and *PL* gene families are absent in *L. bicolor*. Figure below shows a representation of homogalacturonan and the cleavage sites of potential HGMEs.

**Fig. 4:**
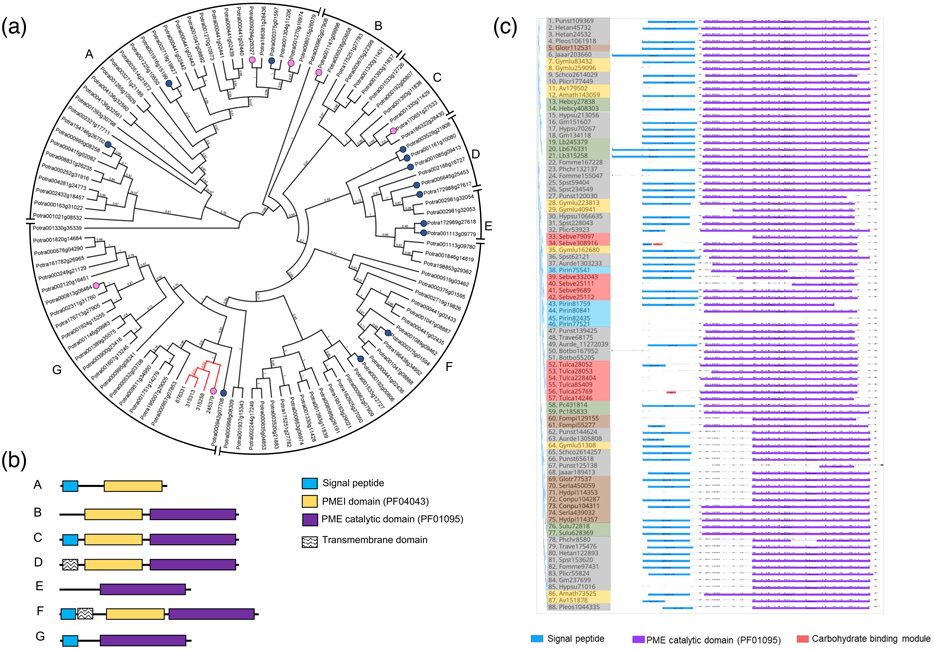
Phylogenetic relationship of PME and PME inhibitor (PMEI) proteins from *P. tremula* (reference genome for *P. tremula* x *P. tremuloides*) and fungi. (a) The unrooted tree was constructed with the maximum likelihood (ML) estimation after alignment of full-length amino acid sequence of 74 *P. tremula* and 4 *L. bicolor* putative gene candidates (red text). Numbers at nodes indicate the percentage of bootstrap scores from 1000 replicates. Clades are classified based on the presence of various domains among most of the members. A graphical representation of these domains is given (b). Gene IDs and sequences of *P. tremula* and *L. bicolor* were extracted from the Plant Genome Integrative Explorer Resource: PlantGenIE.org (Sundell *et al*., 2015) and from the Genome Portal of the Department of Energy Joint Genome Institute (Grigoriev *et al*., 2012). Up-regulated genes in *L. bicolor / P. tremula* x *P. tremuloides* interaction are marked with circles in magenta and down-regulated genes are marked with darkblue circles (a). (c) PME protein domain analysis of various fungal species with different lifestyles. Protein IDs are color-shaded based on fungal lifestyle, namely ectomycorrhizal fungi (green), white-rot fungi (grey), brown-rot fungi (brown), endophytes (blue), orchid mycorrhizal fungi (red), and saprotrophs (yellow). All fungal PMEs lack the PMEI domain and most of them contain signal peptides.

De-esterification by PMEs precedes HG degradation by the activity of HG depolymerizing enzymes like Polygalacturonases (PGs) or Pectin lyases (PNLs) (Sénéchal *et al*., 2014). Our transcriptome profiles showed a significant up-regulation of only one out of 38 *Populus PGs*, but three out of six *L. bicolor PGs* during the interaction. Interestingly, we also noted up-regulation of three *Populus* PG-inhibitors (PGIs), which can potentially influence both plant and fungal PG activity (Federici *et al*., 2001; Protsenko *et al*., 2008; Benedetti *et al*., 2011). Although *PNLs* appear to be absent from the *Populus* genome, one *Populus Pectate lyase (PL)* was up-regulated at 7 DAC and 4 *Populus PLs* were down-regulated at 21 DAC whereas *PLs* are not present in *L. bicolor*. Similarly, only one *Populus Pectin Acetylesterase (PAE)* was up-regulated at 7 DAC and 4 *Populus PAEs* were down-regulated at 21 DAC; this group of enzymes was again not detected in *L. bicolor*. Altogether, our transcriptome analysis revealed ECM-specific regulation of HG modifying gene family members associated with HG de-esterification, deacetylation and depolymerization during *L. bicolor* and *P. tremula* x *P. tremuloides* interactions. Concerning potential HG biosynthesis genes in *Populus*, only one *Galacturonosyltransferase* was up-regulated, the others were down-regulated (Fig. S9). We did not find any differential expression of *Populus* pectin methyl transferases (Fig. S9). Therefore, the data are inconclusive regarding whether pectin synthesis in *Populus* is affected by ECM formation.

Comparing our data to two published datasets of time-course gene-expression profiles during *Populus - L. bicolor* interactions in a soil system (Veneault-Fourrey *et al*., 2014; Plett *et al*., 2015), we noticed an overlap of the differentially expressed *LbCAZymes* in both our and the previous studies (Fig. S10). Out of our 63 *L. bicolor CAZYmes* that were differentially expressed at at least one time-point in our study, 33 (Veneault-Fourrey & Martin, 2011; Veneault-Fourrey *et al*., 2014) and 45 (Plett *et al*., 2015) were respectively found to overlap with our results (Table S7). However, none of the differentially expressed *Populus CAZYmes* (Table S8) from our study was detected in the other studies, probably owing to the use of a different *Populus* hybrid in our study. Interestingly, among the overlapping *LbCAZYmes*, a majority corresponded to genes induced during the late phases of ECM formation in soil systems (clusters C and D in Veneault-Fourrey *et al*. (2014)). This can be explained by the fact that, in soil systems, only few hyphae are in contact with the plant in the early stages, whereas these phases progress much more quickly in *in vitro* systems. The majority of genes (33 out of 45, Fig. S10) that overlapped with LbDEGs in Plett et al. (2015) belonged to what they identified as regulon L_H_ and were those significantly differentially regulated during the aggregation phase and maintained their regulation until time-points of mature ECM in association with *P. trichocarpa* in this report.

### Pectin de-esterification during ECM development

To decipher the spatial and temporal HG methylesterification patterns during ECM formation we used immunofluorescence microscopy with LM19 and LM20 antibodies that bind preferentially to de-esterified HG or highly methylesterified HG, respectively (Verhertbruggen *et al*., 2009). In radial walls between adjacent epidermal cells, which loosens during Hartig Net formation, the abundance of LM19 epitopes was visually higher, and the one of LM20 epitopes was lower in sections through ECM roots (14 DAC) as compared to similar tissues in control roots (Fig. **5a-d**, arrowed). In control roots the LM19 signal was patchy and mostly visible at cell corners and in the tangential wall separating epidermis and cortex cells (Fig. 5a), whereas in ECM sections the signal was strong and homogenous all around the epidermis cells (Fig. 5c). Labelling with LM20, on the other hand, gave a more homogenous signal in the epidermis of control root sections (Fig. 5b) and decreased in the epidermis and the outermost cortex layer of ECM sections to a signal almost solely visible in the cell corners (Fig. 5d). Quantification of the LM19:LM20 ratio confirmed that *in situ* methylesterification levels were significantly modified in ECM compared to control roots. There was a significant increase in the LM19:LM20 ratio from 7 DAC onwards both in the epidermal layer (Fig. **5e**), which is in physical contact with the fungus, and even in underlying root cells (Fig. **5f**). Similar observations were made in successive sections taken at 100 *µ*m intervals between 150 *µ*m and 550 *µ*m from the root tip (Fig. **S11**). This suggests that HG undergoes de-esterification throughout ECM development starting prior to Hartig Net formation, as concluded from the 7 DAC time-point when only a fungal mantle but no Hartig Net is yet present (Fig. **S3**). Transmission electron micrographs (TEM) coupled with immuno-gold labelling confirmed that increased levels of LM19 epitopes were present in the plant cell-wall within the Hartig Net compared to the epidermal wall of the control roots (Fig. S12, quantified in Fig. S13). We observed that LM19 epitopes were mainly limited to the interface of plant and fungal cell-walls and were absent from fungal cell-walls and from the space between adjacent fungal cells in the Hartig Net. The increased abundance of de-esterified HG was accompanied by a significant increased PME activity in 14 DAC colonized roots (with contributions from both plant and fungal PMEs) as compared to control roots or free-living mycelia in two separate experiments (Fig. 6).

**Fig. 5:**
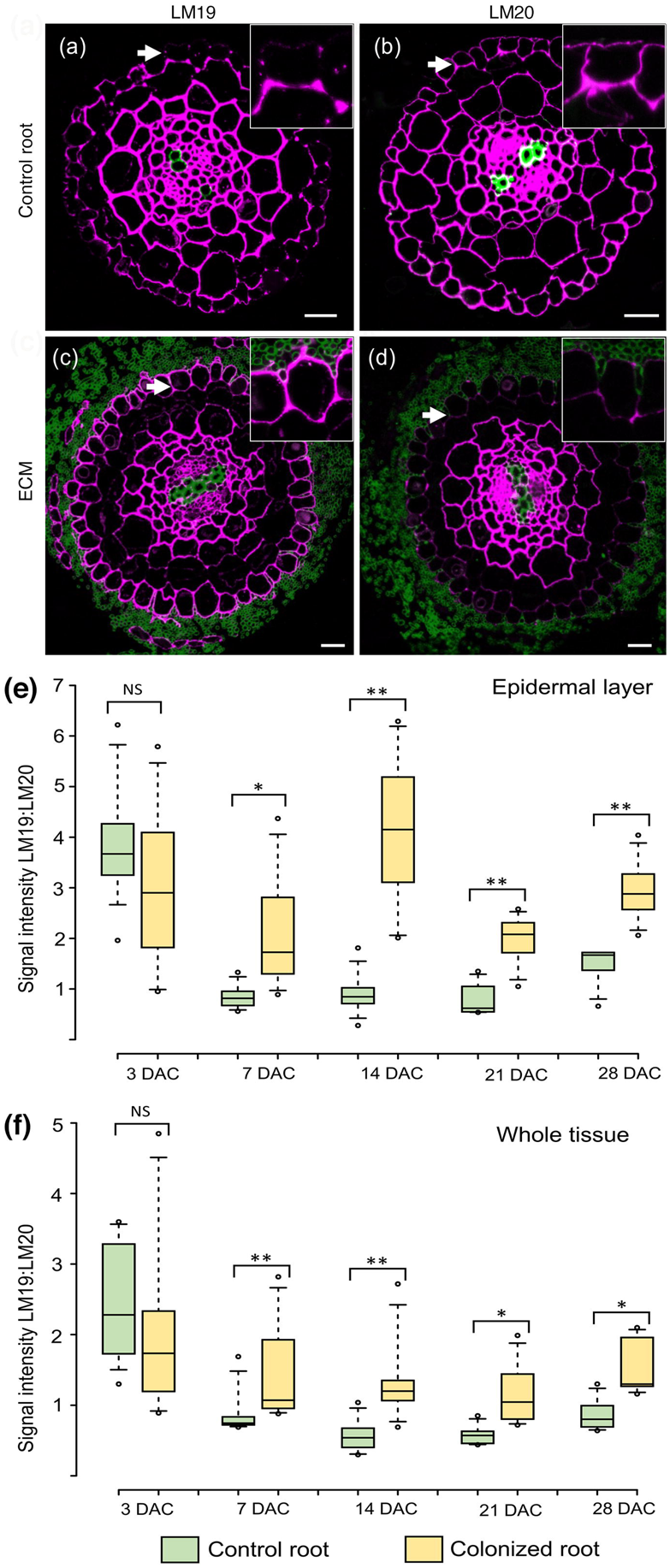
Immunolocalization with pectin antibodies LM19 (recognizing de-esterified homogalacturonan, HG, (a, c)) and LM20 (recognizing methylesterified HG (b, d)) in 1µm thin cross-sections of control roots without fungus (a, b) and colonized roots (c, d) collected at 14 days after contact (DAC) (or control without fungus). *L. bicolor* is visualized by labelling with WGA-AF488 (green); and plant cell-wall homogalacturonan is visualized with LM19 and LM20 antibody conjugated Cy5 (magenta). White arrows indicate the epidermal layer where the difference of antibody labelling abundance is evident between ECM and control roots. Scale bar = 20 µm. Insets in images a-d show a magnified view of three adjacent epidermal cells. Fluorescence intensity ratio of LM19 and LM20 in the epidermal layer (e) and whole tissue area (f), respectively, at the indicated time-points after contact. Data was collected from ten observations per root and four to five roots per timepoint and condition. ** p-value <0.01, * p-value <0.05 with the student t-test.

**Fig. 6.**
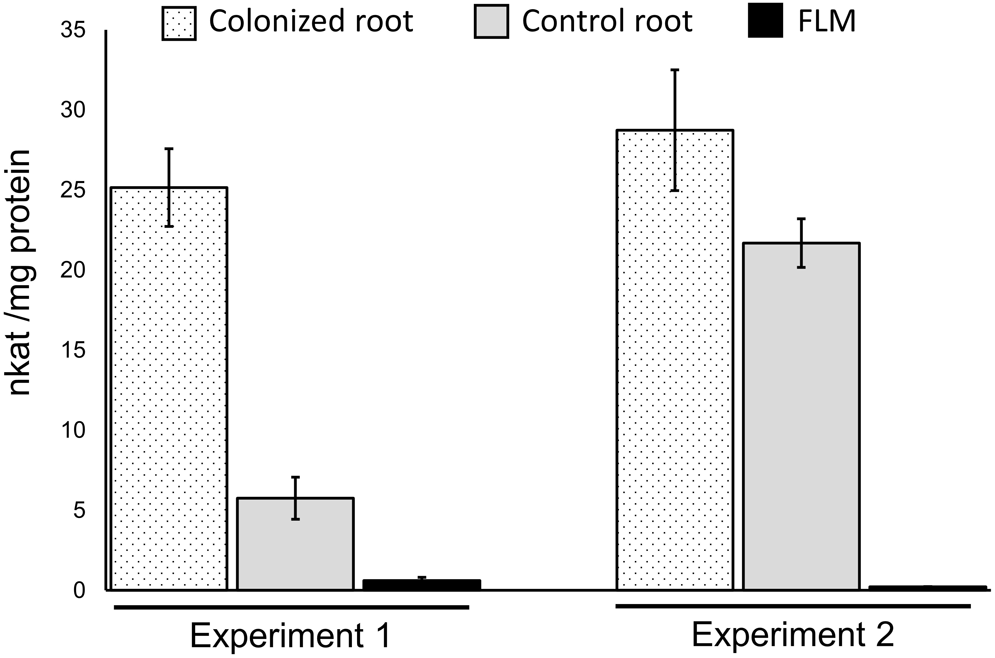
PME activity of pooled lateral roots (50 mg) collected from *P. tremula* x *P. tremuloides*/*L. bicolor* colonized roots at 14 days after contact (DAC) and control roots without fungus from two independent experiments each with three biological replicates. PME activity is significantly increased in ECM tissue in both experiments. Statistical analysis was performed using Student t-test. P value L0.01. Error bars = standard deviation.

### Transgenic alteration of *L. bicolor* PME transcripts alters PME activity level

To assess the function of LbPMEs during ECM formation, we altered *LbPME* transcript levels in *L. bicolor* through RNAi or overexpression constructs driven by the constitutive *Agaricus bisporus gpdII* promoter (Fig. **S1**). For RNAi silencing, the promoter was followed either by a hairpin construct generating siRNAs against *LbPME1* (*LbPME1*_RNAi) or a combined hairpin construct generating siRNAs against *LbPME1* and all three other PMEs that are closely related in sequence (*LbPME1-4*_double RNAi, Figs S1, S14). In free-living mycelia of the three selected *LbPME1-4*_double RNAi lines, the levels of all four *LbPMEs* were significantly reduced by 60% compared to levels in wild-type free living mycelia (Figs **7a**, **S2**). In the three selected single RNAi lines, specifically *LbPME1* expression but not *LbPME2-4* was reduced to a similar extent as in the double_RNAi lines. Finally, in the three selected *LbPME1* over-expressor lines, *LbPME1* transcript levels were increased by 4.9 to 7.9-fold compared to wild-type free-living mycelia, while no effect was observed on transcript levels of the other *LbPMEs* (Figs **7b**, **S2**). We also assessed the expression level of the three ECM-induced *LbPGs* (Fig. S6) in free-living mycelia. *LbPG* JGI ID #388191 was not detected with two different primer pairs, *LbPG2* (JGI ID #612983), and *PbPG1* (JGI ID #613299, *LbGH28A* in Zhang *et al*. (2022)) did not show any significant alteration except for a slight induction of both genes in LbPME1 RNAi line 31 (Fig. S15). Since the transcript sequences or *LbPG1* and *2* are 94% identical we may however observe cross-hybridization of primers, indicated by the similar expression patterns observed across all samples (Fig. S15). In line with *LbPME* expression results, PME activity assays on free-living mycelia grown on P20 supplemented with freeze-dried root cell-wall powder indicated significant reductions of LbPME activity in all *LbPME1*_RNAi and *LbPME1-4*_double RNAi lines (Fig. **7c**). Conversely, two of our three over-expressor lines showed significantly increased LbPME activity. However, there was no strong correlation between transcript levels and PME activity levels inside mycelia samples (Fig. **S16**), which may be due to secretory loss of PME from the mycelium into the growth media. We selected the lines that had both confirmed altered *LbPME* transcript and activity levels for further characterization. To test whether altered PME activity affects the fungal ability to utilize pectin as a carbon source, we grew transgenic lines in modified P20 medium containing pectin that was highly methyesterified or unesterified as the only carbon source. Whilst on P20 (with glucose as carbon source) none of the transgenic lines showed a significant difference in growth compared to control lines, their growth was significantly reduced in both types of pectin containing media irrespective of whether *LbPME* levels had been up- or down-regulated (Fig. **8**). In these pectin media, the hindered growth of PME RNAi lines with restricted PME activity was expected as it suggests that RNAi lines have a reduced ability to utilize pectin. The hampered growth of the over-expressor lines was unexpected.

**Fig. 7.**
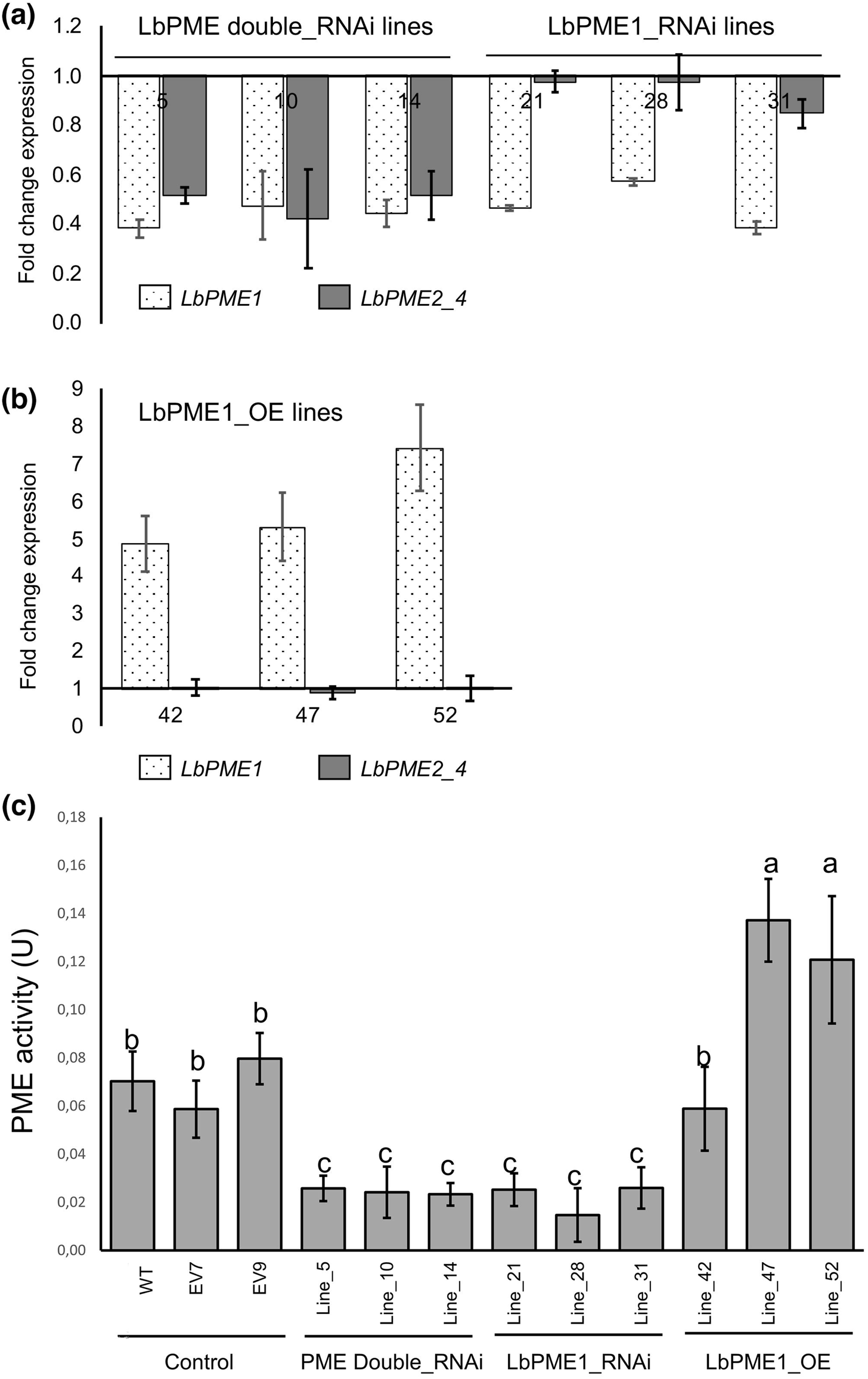
Relative normalized expression of *L. bicolor* PME transcripts in free-living mycelia of the selected transgenic fungal lines (indicated by number) at 14 days of growth compared to wild-type control (a, b). Error bars indicate standard error of mean. PME activity in the free-living mycelia of transgenic *L. bicolor* lines grown on modified P20 medium containing powdered poplar root cell-wall (c). PME activity was measured as the milli-unit when one unit released 1.0 micro-equivalent of acid from pectin per min at pH 7.5 at 30 °C. EV7 and EV9 refer to empty vector mock transformant lines. Statistical analysis was performed using Fisher’s LSD test. The PME activity differences in transgenic fungal lines marked with different letters are statistically significant. Error bars indicate standard deviation.

**Fig. 8.**
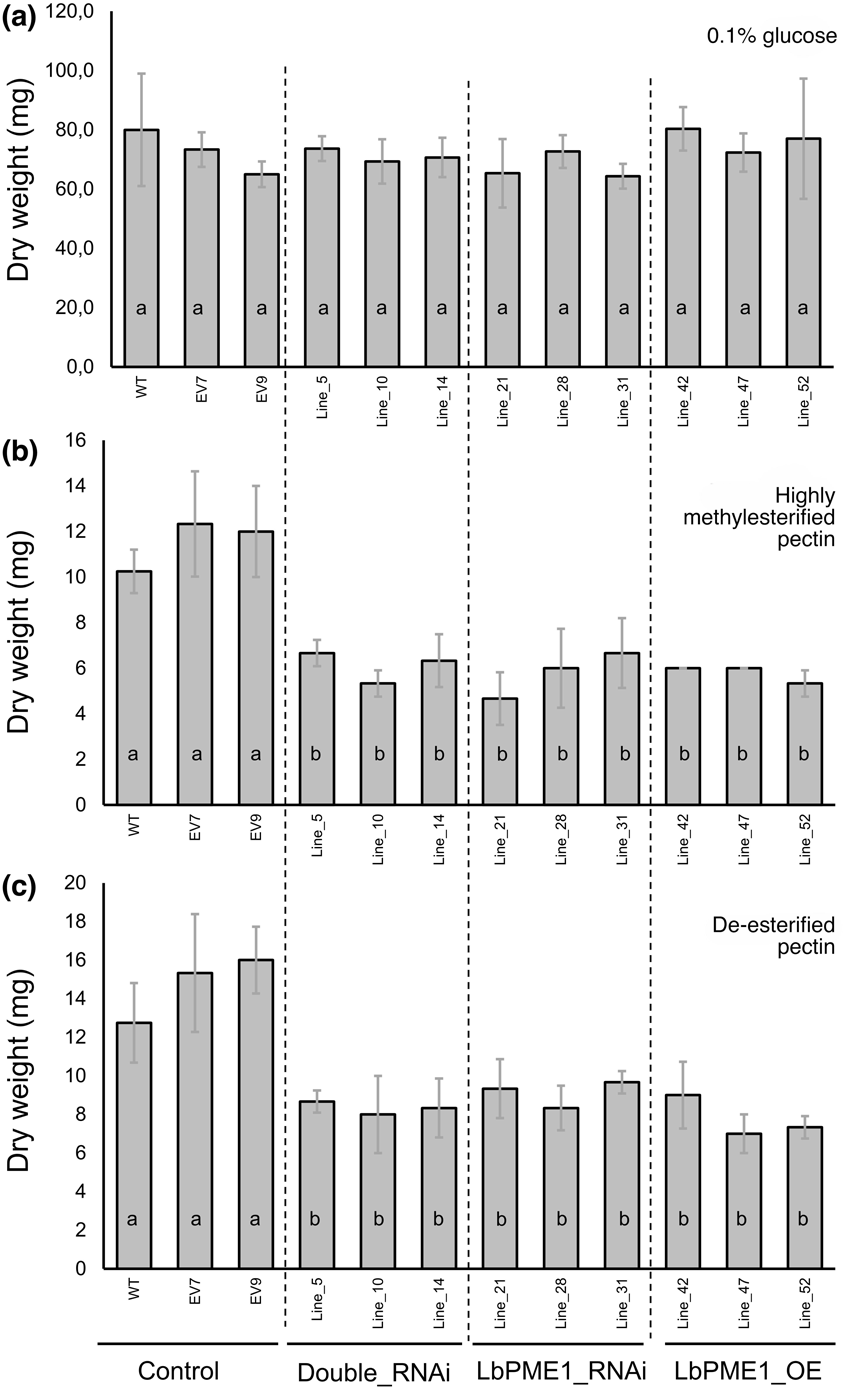
Growth of free-living mycelia of *LbPME* transgenic lines in medium containing various sources of carbon. Medium includes conventional P20 with 0.1% glucose (a), modified P20 with highly methylesterified pectin (b) and with de-esterified pectin (c) as the carbon source. Growth was measured on a dry weight basis. Error bars indicate standard deviation. Significant differences among samples are indicated by unique letters according to a Fisher’s least significant difference (LSD)-ANOVA test (P-value ≤ 0.05).

### Transgenic alteration of *LbPMEs* alters ECM formation and depth of Hartig Net

To assess the role of LbPMEs in ECM development, we performed interaction assays between the selected *L. bicolor* lines and *P. tremula* x *P. tremuloides* and analysed root phenotypes at 14 DAC, which is the earliest time-point when ECM can be well distinguished visually and the Hartig Nets are detected in sectioned material (Fig. S3). Depending on the individual line we found a 22% - 51% decrease in the number of swollen ECM roots with visible mycelia attachment (Fig. **S17**) in interactions with LbPME1 RNAi_lines and LbPME1-4_double RNAi lines (Fig. **9**). Unexpectedly, both selected LbPME1 over-expressor lines showed reduced ECM frequency comparable to the RNAi lines.

**Fig. 9.**
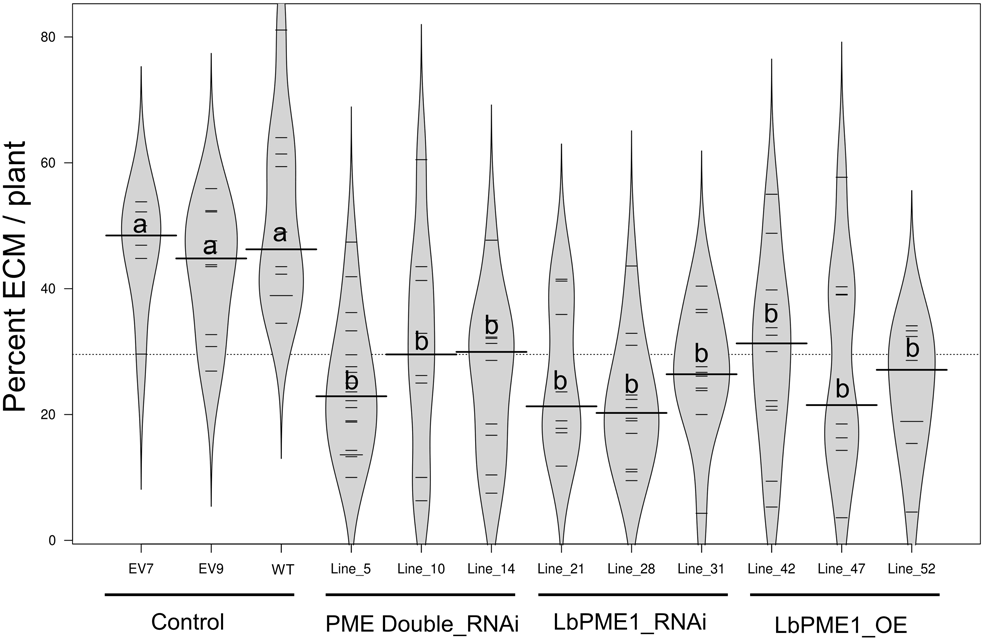
Bean plot showing the frequency of *P. tremula* x *P. tremuloides* ECM at 14 days after contact (DAC) with wild-type or genetically modified *L. bicolor* lines. The width of each column indicates the density of observations and the short horizontal lines represent ectomycorrhiza (ECM) number per observed plant. Long horizontal lines indicate the mean value. Significant differences among samples are indicated by unique letters according to a Fisher’s least significant difference (LSD)-ANOVA test (P-value ≤ 0.05).

To confirm the results from morphological observations, we assessed the expression of symbiosis-associated marker genes - two previously characterized from *L. bicolor* and one from *Populus -* in RNA from randomly collected lateral roots at 14 DAC (containing non-colonized, colonized and ECM roots) in interactions with the transgenic *L. bicolor* lines: *PF6.2* (JGI ID # 469385) (Kim *et al*., 1998; Podila *et al*., 2002; Hiremath *et al*., 2013), *Aquaporin* (JGI ID # 671860) (Dietz *et al*., 2011; Kohler *et al*., 2015) and the sugar transporter *PtSWEET1* (Neb *et al*., 2017). We confirmed that these genes are induced in co-cultures with control (empty vector) *L. bicolor* lines compared to conditions where both organisms were grown separately (Fig. **10a, b**). Consistent with morphological observations, *PF6.2, Aquaporin* and *PtSWEET1* were significantly down-regulated in co-cultures with the RNAi and over-expressor lines compared to co-cultures with empty vector control lines at 14 DAC (**Fig. 10c-e**). With this experiment we also confirmed that *LbPME1* transcript levels in all selected over-expressor and RNAi lines were altered similarly in free-living mycelium (Fig. **7a**) and in mycelium in contact with roots (Fig. **10f**). Finally, we examined expression levels of ECM-induced *LbPGs* (using the primer for JGI ID # 612983 that likely also recognizes ID #613299, LbGH28A) as changes in *PME* expression levels may affect the expression of the HG-degrading *PGs* (Hadfield & Bennett, 1998). Although there was no alteration of *PG* expression in the free-living mycelia of PME altered lines (Fig. S15), we found a significant reduction of *LbPG* transcript levels in co-cultures with RNAi and OE lines compared to co-cultures with control *L. bicolor* lines (Fig. **10g**). This suggests that PG-mediated HG depolymerization may be reduced in co-cultures with RNAi and OE lines.

**Fig. 10.**
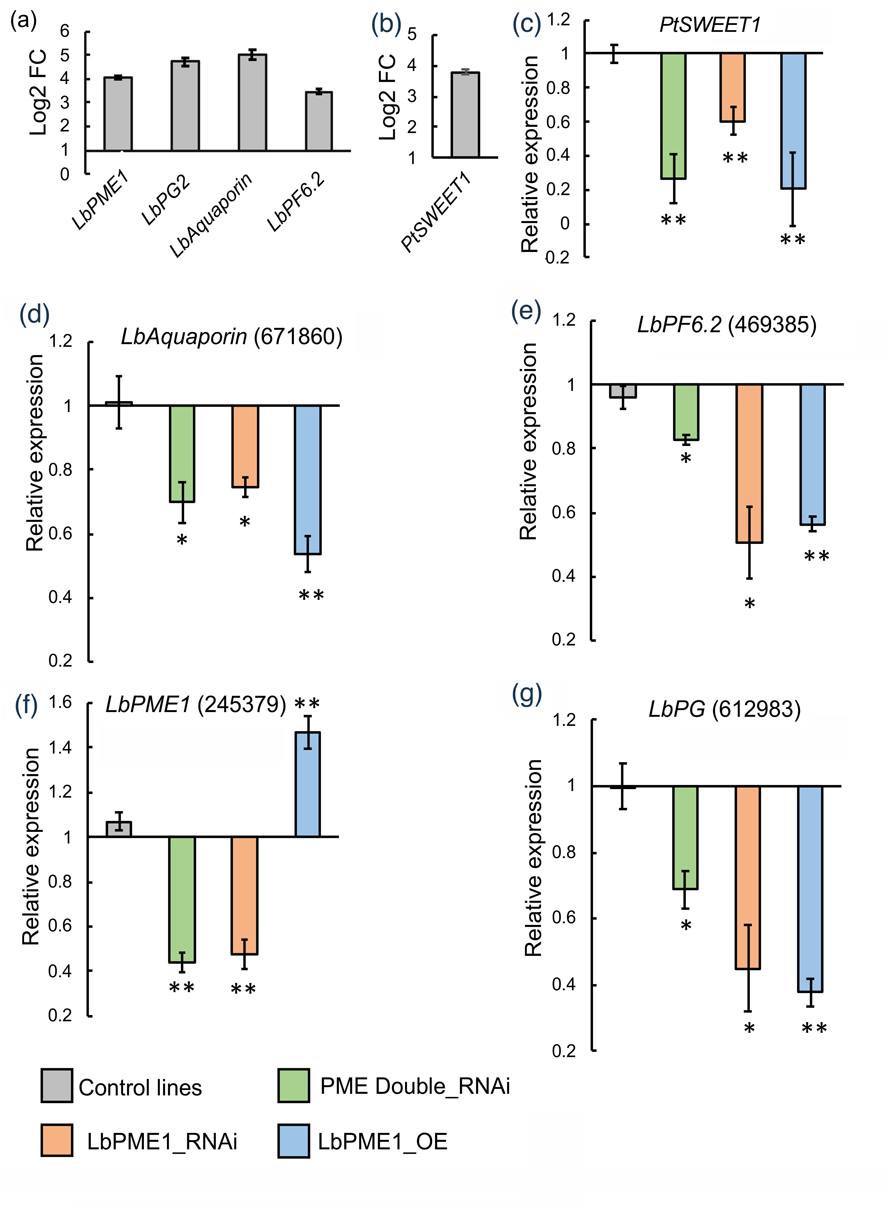
Relative normalized expression of selected homogalacturonan modifying genes (HGMEs) and ectomycorrhiza-related genes of *L. bicolor* and *Populus* in co-culture tissue (comprising ECM, colonized roots and non-colonized roots) at 14 DAC determined by qPCR. Log_2_ fold induction of the gene of interests in control *L. bicolor* (average of wild-type and mock vector control lines) interacting with *P. tremula* x *P. tremuloides* (14 days after contact, DAC) relative to free-living mycelia without the plant (a). Fold-change of genes of interest in co-cultures with the transgenic lines relative to co-cultures with control lines of *L. bicolor* (b-e). For each construct, co-culture samples with the three selected transgenic *L. bicolor* lines were considered as three biological replicates. Expression of each gene was normalized against their expression in wild-type *L. bicolor* interacting with *P. tremula* x *P. tremuloides*. Student t-test: ** indicates P<0.01, * indicates P <0.05 compared to wild-type. Error bars indicate standard error of the mean.

We furthermore measured the depths and areas of Hartig Nets in cross-sections of ECM with the RNAi and OE lines compared to ECM with wild-type *L. bicolor*. As micrographs revealed, Hartig Net depths in ECM with the RNAi lines were reduced by 60 % compared to ECM with wild-type fungus (Figs **11a**, Fig. **S18c-d**). This suggests that the reduced PME activity restricts the ability of the RNAi lines to loosen plant cell-walls and to induce root cell separation by the RNAi lines. ECM with *LbPME1* OE lines, on the contrary, produced slightly deeper Hartig Nets (on average 18% deeper than wild-type) (Figs **11a**, **S18e**). The expression of *LbPMEs* expression also affected the Hartig Net area in between adjacent root cells representing the colonized area in the apoplastic region (Fig. **11b**). Depending on the lines, the area was reduced by about 40% among the RNAi lines and increased by about 40% among the OE lines. Using LM19 antibodies to detect de-esterified HG in ECM cross-sections at 14 DAC, we detected reduced labelling in radial epidermal walls of ECM with RNAi lines (significant at p<0.05 for LbPME double RNAi lines, visible trend for LbPME1 single RNAi lines (p=0.067)) as compared to ECM with control lines or LbPME OE lines, despite variation between the individual lines (Fig. S18f). Taken together, these results show that *L. bicolor* requires an optimal PME level to form ECM, whereas suboptimal LbPME expression, whether reduced or increased, negatively affected ECM frequency. While transgenic reduction of LbPME expression lowered cell wall de-esterification, Hartig Net depth and ECM frequency, increased LbPME levels did not increase ECM frequency nor de-esterification in the epidermal layer, but resulted in a deeper Hartig Net.

**Fig 11.**
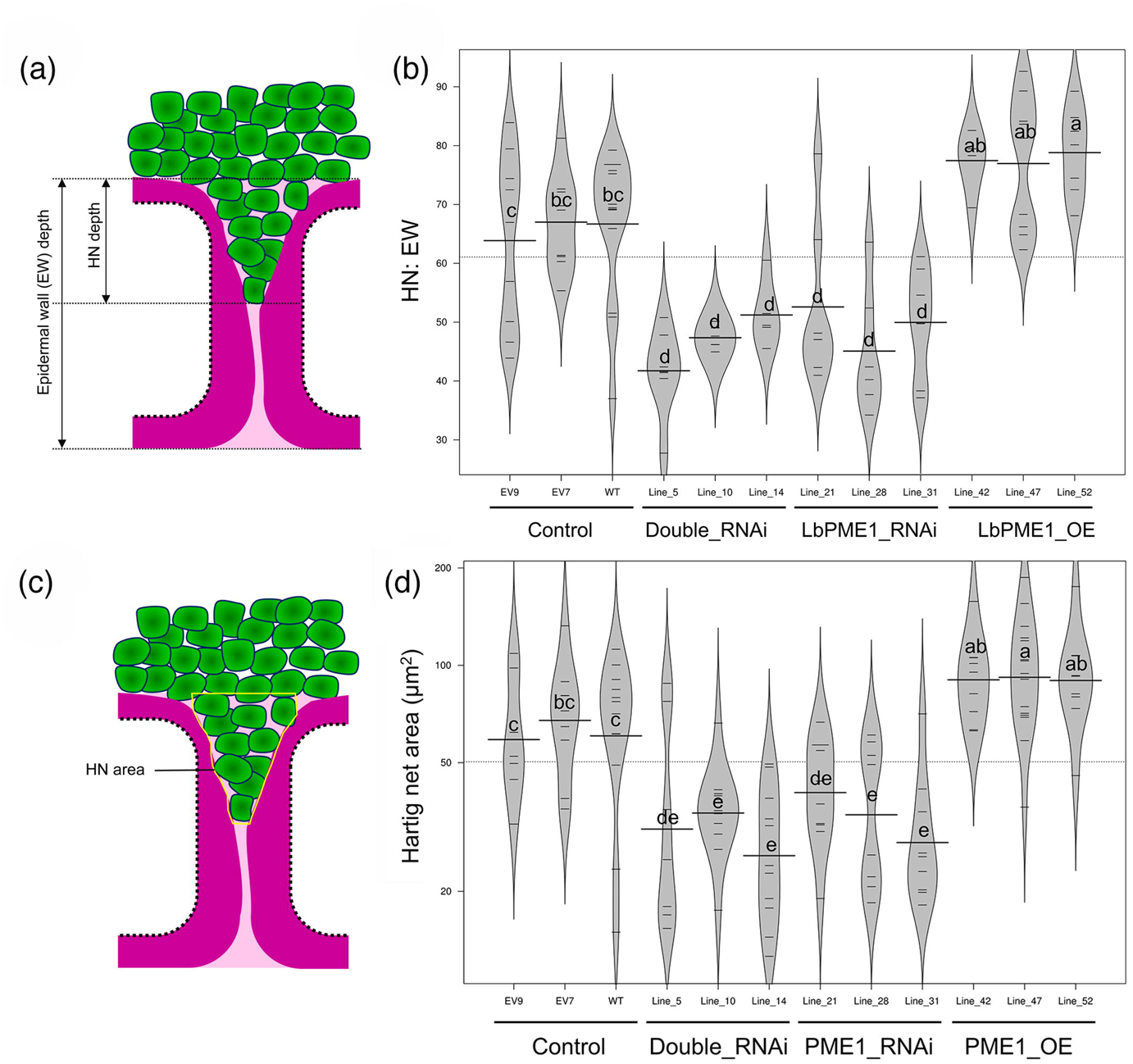
Hartig Net depth and area in *P. tremula x P. tremuloides* ECM at 14 days after contact (DAC) with wild-type or genetically modified *L. bicolor*. Hartig Net depth is represented as the percentage of the area of the radial epidermal wall covered by *L. bicolor* (a). Bean plot showing Hartig Net depth as the ratio of Hartig Net and epidermal wall among various interactions (b). Bean plot showing the ratio of Hartig Net and epidermal wall among various interactions (c). While long horizontal lines indicate the mean value, small horizontal lines represent individual data points and polygons represent the estimated density of the data (b, c). Significant differences among lines are indicated by unique letters according to a Fisher’s least significant difference (LSD)-ANOVA test (P-value ≤ 0.05).

## Discussion

The molecular mechanisms of cell-wall remodelling associated with ECM development are just beginning to be understood despite their importance for Hartig Net formation (Balestrini & Bonfante, 2014). As revealed by whole genome sequences, most ECM fungi have a reduced set of *PCWDEs* compared to their saprotrophic ancestors. The loss of *PCWDEs* has been suggested to be the result of potential mechanisms to avoid pattern triggered immunity responses from host plants and to make colonization possible . Nevertheless, from the transcriptome analyses of various ECM interactions, including our present, it is evident that ECM-specific induction of *PCWDEs* commonly occurs, and is likely to be necessary for ECM formation (Sebastiana *et al*., 2014; Veneault-Fourrey *et al*., 2014; Kohler *et al*., 2015). A recent study has shown that ECM formation requires the *L. bicolor* endoglucanase, LbGH5LCBM, acting on plant cell-wall cellulose. Silencing of this gene reduced *L. bicolor’s* ability to form ECM (Zhang *et al*., 2018). Like cellulose, HG is a major component of the plant cell-wall, although it is mainly localized in the middle lamella. HG is considered a key polymer for cell-to-cell adhesion (Bouton *et al*., 2002; Daher & Braybrook, 2015) and our study, together with another recent report (Zhang *et al*., 2022), reveal that modulation of the HG-rich middle lamella is a prerequisite for complete Hartig Net formation and entails the action of members of two families of *L. bicolor* HG modifying enzymes (HGMEs).

The increased abundance of de-esterified HG especially in the area where the Hartig Net actively forms from 7 DAC onwards (Fig. **5**), is in line with a previous study that, using JIM5 antibody labelling, demonstrated the presence of de-esterified HG epitopes in the proximity of the Hartig Net in *Tuber melanosporum* and *Corylus avellana* ECM (Sillo *et al*., 2016). However, Sillo et al. (2016) reported also that, using a carbohydrate microarray, all HG epitopes along with other PCW polysaccharides significantly decreased in ECM with *T. melanosporum* as compared to non-colonized root tissues of *C. avellana.* This discrepancy is most probably due to the different nature of the materials used in this study. Only the plant partner contains HG; thus, when bulk material from roots and ECM (root and fungus) are extracted and compared, and the results from ECM are normalized on total ECM weight, one can expect to detect a reduced (diluted) amount of HG in extracts from ECM roots, regardless of the methylesterification state *per se*. Therefore, we have taken advantage of quantitative immuno-fluorescence microscopy to assess the HG methylesterification state at a tissue level. To normalize for root-to-root variation of HG methylesterification, we have compared roots and ECM of similar age at specific time-points, by measuring the signal intensity ratio arising from LM19 and LM20 epitopes (Fig. **5**). From this we can deduce the proportion of de-esterified and highly methylesterified HG in the ECM composite material. The HG de-esterification pattern during the interaction of *Populus* with *L. bicolor* resembles several other plant interactions with fungal root pathogens, i.e. pectin gets de-esterified by PMEs to make tissues more vulnerable to pathogen CWDEs (Lionetti *et al*., 2012; Ma *et al*., 2013; Fan *et al*., 2017). Our finding of increased HG de-esterification during Hartig Net formation is consistent with our further observation that PME activity is increased in ECM roots. Even though it is not possible to conclude from this experiment whether it is the plant or the fungal partner or both that exert the enhanced activity in ECM tissues, our further analyses support the importance of fungal LbPMEs in the process.

PMEs are the key enzymes controlling the HG methylesterification status, whereas galacturonosyl transferase or pectin methyltransferase activity does not affect the methylesterification degree of HG (Mouille *et al*., 2007; Wolf *et al*., 2009). However, given our current state of knowledge, we only partly understand the action and consequences of PME-mediated HG de-esterification. It may either lead to cell-wall loosening by PGs and PLs, or it may cause cell-wall rigidification through calcium cross-linking and the formation of the so-called egg-box (Sénéchal *et al*., 2014; Hocq *et al*., 2017; Wormit & Usadel, 2018). Specific physiological conditions may determine this fate. We assume that PME-mediated HG de-esterification throughout ECM development leads rather to HG degradation and cell-wall loosening. This argument is supported by a significant induction of three fungal *PGs* and one plant *PG* potentially catalysing HG depolymerization in ECM, and advocates for HG breakdown during ECM formation. Furthermore, recent evidence has proved that the fungal polygalacturonase LbGH28A indeed acts on pectin (Zhang *et al*., 2022). LbGH28A could be a likely actor downstream of the action of LbPME, where LbPME1 catalyzes HG de-esterification and LbGH28A breaks down the de-esterified HG. As *L. bicolor* only possesses these two types of pectin related enzyme families (Veneault-Fourrey *et al*., 2014) further catalytic functions may be carried out by plant enzymes (Figs 12, S6) during the earlier stages of ECM formation. Three plant *PMEs* were up-regulated up to the 14 DAC time-point but thereafter, we only observed down-regulation of plant PMEs (Fig. **S6**). The product of the two plant *PMEIs* induced at 21 DAC may inhibit any remaining expressed plant PMEs (Juge, 2006; Pelloux *et al*., 2007; Sénéchal *et al*., 2014; Wormit & Usadel, 2018). Plant PMEIs do not, however, seem to be capable of inhibiting fungal and bacterial PME activity due to the absence of critical residues for PME-PMEI interactions in microbial PMEs (Di Matteo *et al*., 2005; Lionetti *et al*., 2007; Reca *et al*., 2012) including *LbPMEs* (Fig. **4**). The significant up-regulation of *LbPME1* throughout the different stages of ECM development, that precedes detectable de-esterification (from 7 DAC), and its potential insensitivity to PMEI-mediated inhibition led us to consider this gene as a potential key contributor to HG de-esterification. *LbPME1* is also the only PME induced in interaction with other tree species, namely *Populus trichocarpa* and *Pseudotsuga menziesii* (Plett *et al*., 2015). This suggests that *LbPME1* is an ECM-associated fungal PME isoform potentially regulating cell-wall remodelling during Hartig Net formation.

**Fig 12.**
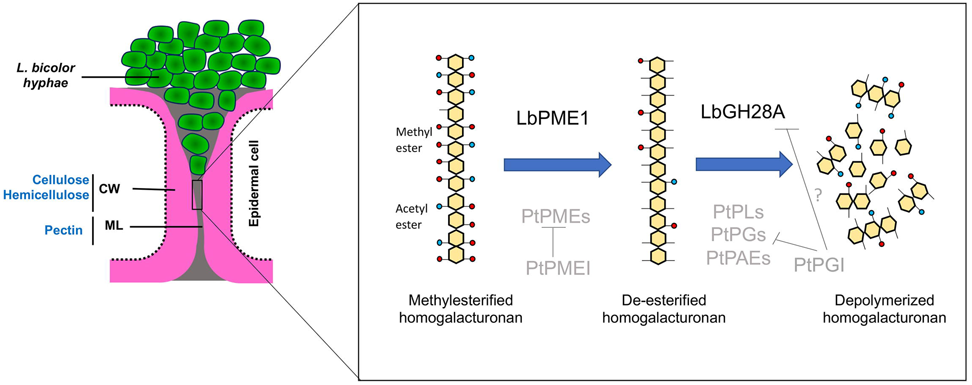
Model of pectin modification during Hartig Net formation *in L. bicolor / Populus* interaction. Fungal players are indicated in black and putative plant players in grey. Homogalacturonan (HG) is de-esterified with the help of LbPME1 as shown in this study. De-esterified HG is sensitive to degradation catalyzed by enzymes of the LbGH28 family (Zhang *et al*., 2022). Further plant enzymes may contribute to this process in the early phase of root colonization as indicated by our transcriptome data (Fig. S6). Plant PMEIs are unlikely to act on fungal LbPME due to the lack of a PMEI binding site in LbPME, PGIs may act on plant and fungal enzymes. PME = Pectin methylesterase, PMEI = Pectin methylesterase inhibitor, PAE = Pectin acetylesterase, PL = Pectate lyase, PG = Polygalacturonase, PGI= Polygalacturonase inhibitor, CW = Cell Wall, ML = middle lamella

Our functional approach using single and double LbPME RNAi lines shows that the ECM-induced *LbPME1* is not functionally redundant to *LbPME2* to *4*, i.e. suppression of *LbPME1* did not induce other *LbPME* members (Fig. **7a, b**), nor did other LbPMEs compensate functionally for *LbPME* repression. The latter was supported by our finding that PME activitiy of single RNAi and double RNAi lines did not significantly differ (Fig. **7c**), and that the double RNAi construct did not cause additional phenotypes as compared to the single RNAi construct (Figs **9**, **11**). The fact that LM19 signals were reduced at 95% significance level only in the double RNAi but not the single RNAi lines despite a visible trend, can be attributed to variations between the individual lines (Fig. 18f). As we observed the expected increased PME activity in *L. bicolor* lines overexpressing LbPME1 and decreased PME activity in RNAi lines (Fig. **7c**), we deemed *in vitro* enzyme characterization of *LbPME1* unnecessary.

The fact that RNAi lines had a lower ECM frequency, reduced Hartig Net depth, and a decreased expression level of marker genes, suggests that HG de-esterification, with the contribution of *L. bicolor LbPME1*, is needed for full Hartig Net formation. This also supports the idea that this step is a prerequisite for enzymatic breakdown of HG polymers by PGs and PLs (Verlent *et al*., 2005), with potential contribution by LbGH28 (Zhang *et al*., 2022). Strikingly, although OE lines had significantly increased Hartig Net depth and area (Fig. **11**) in roots that managed to form ECM with this line, overall ECM frequency was decreased in the same way as in RNAi lines, as was marker gene expression. Furthermore, the expression of an ECM-induced polygalacturonase gene, *LbPG2,* was significantly reduced in colonized roots with all transgenic lines when compared to roots colonized by wild-type fungus (Fig. **10**), which indicates a correlation with reduced HG breakdown and ECM formation. One could hypothesize that the overproduction of LbPME1 in the OE lines leads to blockwise de-esterification of HG, which creates potential binding sites for Ca^2+^ and egg-box formation (Cabrera *et al*., 2008), thereby stiffening the middle lamella and reducing the access of depolymerizing enzymes like PGs and PLs for HG breakdown (Ralet *et al*., 2001; Pelloux *et al*., 2007; Ngouémazong *et al*., 2012; Sénéchal *et al*., 2014). However, we estimate this scenario rather unlikely as the existence of egg-box structures *in planta* has recently been questioned and atomic force microscopy suggests that overexpression of PME results in a reduced stiffness of cell-walls (Hocq *et al*., 2017). Nevertheless, such structures may have formed in the growth experiments of free-living mycelia on plates with pectin as a carbon source (Fig. **8**). Under those conditions, it is possible that an over-production of LbPME by the OE lines leads to block-wise de-esterification of HG which acts as a nucleation site of ions like Mg^2+^ abundant in the growth medium. Formation of such metal ion bridges may hinder degradation of HG by the activity of PGs (Sénéchal *et al*., 2014) and thereby reduce availability of carbon from the pectin for fungal nutrition. This would then result in reduced growth even in the over-expressor lines, as observed and expected in RNAi lines that have reduced capacity to make carbon available from HG present in the media.

Concerning the *in planta* scenario during ECM formation, which would explain reduced ECM frequency in plants in contact with *LbPME1* overexpressor lines, it is likely that PME overproduction triggers excessive HG degradation involving PGs or PLs. This would then result in an enhanced release of oligogalacturonides (OGs) triggering OG-induced systemic defence responses that could counteract ECM formation. In support of this, several studies provide evidence that OGs, upon binding to wall-associated kinases (WAKs), elicit defence responses, including induction of pathogenesis-related proteins and reactive oxygen species (Anderson *et al*., 2001; Decreux & Messiaen, 2005; Ferrari *et al*., 2013; Benedetti *et al*., 2015). During plant invasion by pathogens, binding of OGs to WAKs triggers MAPK-mediated activation of OG-specific defence gene expression independent of salicylic acid-, jasmonic acid- and ethylene-mediated signalling pathway (Ferrari *et al*., 2007; Chassot *et al*., 2008). Even though overall ECM formation with the overexpressor lines may be restricted by such a systemic *in planta* defence response, at a few positions where ECM manage to form, the enhanced PME activity would lead to deeper Hartig Net formation. Even though we did not detect a higher abundance of LM19 epitopes in radial epidermal walls in ECM with LbPME1 over-expressors at the 14 DAC time-point (Fig. S18f), the deeper Hartig Net and larger Hartig Net area could indicate a quicker progression of Hartig Net formation resulting in a more advanced stage at the timepoint of observation (Fig S3). Such a scenario would go in line with a more strongly triggered defence response in roots as hypothesized above. This hypothesis and the signalling pathways behind it will need to be verified experimentally in the future.

In conclusion, HG de-esterification of the host cell-wall is a critical step for cell-wall remodelling and Hartig Net formation during the *L. bicolor* and *P. tremula* x *P. tremuloides* interaction. The fungal pectin methylesterase *LbPME1* is essential in this process. LbPME1 may play a similar role for the interactions of *L. bicolor* with other *Populus* species and *Pseudotsuga menziesii* because of their expression similarities (Plett *et al*., 2015). Even though ECM fungi may have retained only a small repertoire of pectin-related CAZYmes (Miyauchi et al., 2020, Kohler *et al*., 2015; Martin *et al*., 2016), the reduced machinery in *L. bicolor* makes a significant contribution to ECM formation. Despite a wide retention of the machinery in fungi with different lifestyles, we know a few ECM fungi that lack CE8 (PME) and/or GH28 genes (Kohler et al. 2015, Miyauchi et al. 2020). It will be interesting to explore the mechanisms of pectin remodelling in relationships with fungi that lack the here described mechanism in future studies.

## Acknowledgements

JC was supported by the Kempe Foundations (SMK-1533) and by Formas – a Swedish Research Council for Sustainable Development (942 2105-539). Funding to MK and AGP was provided by grants from National University of Quilmes (UNQ), National Council of Scientific and Technical Research (CONICET), and National Agency for Scientifc and Technological Promotion (ANPCyT), Argentina. We would like to acknowledge support from Science for Life Laboratory, the National Genomics Infrastructure funded by the Swedish Research Council, and Uppsala Multidisciplinary Center for Advanced Computational Science for assistance with massively parallel sequencing and access to the UPPMAX computational infrastructure. The authors acknowledge the facilities and technical assistance of the Biopolymer Analytical Platform (BAP) at SLU/KBC, Umeå University and the Umeå Core Facility Electron Microscopy (UCEM) at the Chemical Biological Center (KBC), Umeå University.

## Author contribution

JC and JFL designed the study, JC conducted and analyzed the experiments. MK designed constructs for transgenic lines used in this project, and prepared and transformed *L. bicolor* under AGP’s supervision. ND and IS performed bioinformatics analysis. JZ carried out insert number and site characterization of transgenic *L. bicolor* lines. JT was involved in setting up PME activity assays. JLF supervised the study. JC and JLF wrote the manuscript with contribution of all authors.

## Data availability

The raw transcriptomics data are available from the European Nucleotide Archive (ENA, https://ebi.ac.uk/ena) under the accession number PRJEB41173.

A GIT-Hub repository of data analysis performed is available under DOI10.5281/zenodo.4629643 at https://doi.org/10.5281/zenodo.4629643 while the code is available at https://github.com/nicolasDelhomme/laccariaBicolorEcmDev/tree/v1.0

## Supporting information

**Fig. S1** Schematic diagram of vector constructs used in this study for altering *L. bicolor* PME expression.

**Fig. S2** qPCR screening of *L. bicolor* RNAi and over-expressor (OE) transformant lines for *PME* expression in free-living mycelium.

**Fig. S3** Gradual development of *L. bicolor* colonization while interacting with *P. tremula* x *P. tremuloides*.

**Fig. S4** Venn diagrams of differentially expressed genes (DEGs) in mycelium in ECM compared to free-living mycelium.

**Fig. S5** Proportion of differentially expressed genes and unchanged genes associated with specific polysaccharide degrading enzymes in *P. tremula* x *P. tremuloides* and *L. bicolor*.

**Fig. S6** Time-course RNA-Seq expression profile of the potential candidates of HG synthesis and modifying enzymes (HGMEs) during *L. bicolor / P. tremula* x *P. tremuloides* interaction.

**Fig. S7** Comparison of gene expression values obtained by RT-qPCR and RNA-seq.

**Fig. S8** Time-course RNA-Seq expression profile of four *L. bicolor* PMEs in colonized roots (ECM) and free-living mycelia (FLM).

**Fig. S9** Differential expression of *P. tremula x P. tremuloides* HG biosynthesis genes during interaction with *L. bicolor* based on RNASeq data from our study.

**Fig. S10** Comparison of differentially expressed Lb CAZymes in our study to those in Veneault- Fourrey *et al*. 2014 and Plett *et al*. 2015

**Fig. S11** Immunolocalization with pectin antibodies LM19 and LM20 at various distances from the root tips.

**Fig. S12** Transmission electron micrographs of immuno-gold labeling of homogalacturonan epitopes present in the epidermis cell wall of ECM and control root cross-sections.

**Fig. S13** Quantification of gold particles in the Hartig Net plant cell-wall and corresponding cell- wall areas of control roots on TEM image.

**Fig. S14** Multiple sequence alignment and sequence identity of *L. bicolor* PME proteins.

**Fig. S15** Relative normalized expression of *LbPG* in free-living mycelium of the PME transgenic lines by qPCR.

**Fig. S16** Correlation of *PME1* transcript level and PME enzymatic activity level in the free- living mycelia of the transgenic fungal lines.

**Fig. S17** Examples of control roots and swollen root tips with attachment of fungal mycelia on colonized roots.

**Fig. S18** Cross-section of Ectomycorrhizal root tips developed by *P. tremula x P. tremuloides* and PME transgenic lines showing the depth of the Hartig Net.

**Table S1** List of primer sequences used in this study. **(Separate excel file)**

**Table S2** Genomic T-DNA integration number and integration sites in *L. bicolor* transformants used in this study. **(Separate excel file)**

**Table S3** Gene expression data of *L. bicolor* (based on JGI genome v2.0 annotation) ectomycorrhiza versus free-living mycelium. **(Separate excel file)**

**Table S4** Gene expression data from *P. tremula* x *P. tremuloides* annotated based on the *P. tremula* genome, ectomycorrhiza versus control root. **(Separate excel file)**

**Table S5** *L. bicolor* genes specifically differentially expressed at only one of the time-points during the interaction (p-adj < 0.01; log_2_ fold change > 0.5) **(Separate excel file)**

**Table S6** *Populus* genes specifically differentially expressed at only one of the time-points during the interaction (p-adj < 0.01; log_2_ fold change > 0.5). **(Separate excel file)**

**Table S7** Differentially expressed (p-adj < 0.01; log_2_ fold change > 0.5) *L. bicolor* CAZYmes in our study compared to literature studies by Veneault-Fourrey *et al*. 2014 and Plett *et al*. 2015 (**Separate excel file**)

**Table S8** Differentially expressed (p-adj < 0.01; log_2_ fold change > 0.5) *P. tremula x P. tremuloides* CAZYmes in our study (**Separate excel file**)

**Methods S1** Bioinformatics

**Methods S2:** PME activity assay in free living mycelium

**Methods S3** Immunofluorescence localization of pectin antibodies in poplar roots

**Methods S4** Cloning, *L. bicolor* transformation and fungal selection

**Methods S5** Characterization of T-DNA insertion number in transgenic *L. bicolor* lines by ddPCR

**Methods S6** Characterization of T-DNA insertion sites in transgenic *L. bicolor* by Plasmid rescue and TAIL-PCR

**Methods S7** Characterization of T-DNA insertion number in transgenic L. bicolor lines by ddPCR

