## Supporting Figure S1-S18, Supporting Methods S1-S7 for "*Laccaria bicolor* pectin methylesterases are involved in ectomycorrhiza development with Populus tremula x Populus tremuloides"

**The following Supporting Information is available for this article:**

**Fig. S1** Schematic diagram of vector constructs used in this study for altering *L. bicolor* PME expression.

**Fig. S2** qPCR screening of *L. bicolor* RNAi and over-expressor (OE) transformant lines for *PME* expression in free-living mycelium.

**Fig. S3** Gradual development of *L. bicolor* colonization while interacting with *P. tremula* x *P. tremuloides.*

**Fig. S4** Venn diagrams of differentially expressed genes (DEGs) in mycelium in ECM compared to free-living mycelium.

**Fig. S5** Proportion of differentially expressed genes and unchanged genes associated with specific polysaccharide degrading enzymes in *P. tremula* x *P. tremuloides* and *L. bicolor*.

**Fig. S6** Time-course RNA-Seq expression profile of the potential candidates of HG synthesis and modifying enzymes (HGMEs) during *L. bicolor / P. tremula* x *P. tremuloides* interaction.

**Fig. S7** Comparison of gene expression values obtained by RT-qPCR and RNA-seq.

**Fig. S8** Time-course RNA-Seq expression profile of four *L. bicolor* PMEs in colonized roots (ECM) and free-living mycelia (FLM).

**Fig. S9** Differential expression of *P. tremula x P. tremuloides* HG biosynthesis genes during interaction with *L. bicolor* based on RNASeq data from our study.

**Fig. S10** Comparison of differentially expressed Lb CAZymes in our study to those in Veneault-Fourrey *et al*. 2014 and Plett *et al*. 2015

**Fig. S11** Immunolocalization with pectin antibodies LM19 and LM20 at various distances from the root tips.

**Fig. S12** Transmission electron micrographs of immuno-gold labeling of homogalacturonan epitopes present in the epidermis cell wall of ECM and control root cross-sections.

**Fig. S13** Quantification of gold particles in the Hartig Net plant cell-wall and corresponding cell-wall areas of control roots on TEM image.

**Fig. S14** Multiple sequence alignment and sequence identity of *L. bicolor* PME proteins.

**Fig. S15** Relative normalized expression of *LbPG* in free-living mycelium of the PME transgenic lines by qPCR.

**Fig. S16** Correlation of *PME1* transcript level and PME enzymatic activity level in the free-living mycelia of the transgenic fungal lines.

**Fig. S17** Examples of control roots and swollen root tips with attachment of fungal mycelia on colonized roots.

**Fig. S18** Cross-section of ectomycorrhizal root tips developed by *P. tremula x P. tremuloides* and PME transgenic lines showing the depth of the Hartig Net.

**Table S1** List of primer sequences used in this study. **(Separate excel file)**

**Table S2** Genomic T-DNA integration number and integration sites in *L. bicolor* transformants used in this study. **(Separate excel file)**

**Table S3** Gene expression data of *L. bicolor* (based on JGI genome v2.0 annotation) ectomycorrhiza versus free-living mycelium. **(Separate excel file)**

**Table S4** Gene expression data from *P. tremula* x *P. tremuloides* annotated based on the *P. tremula* genome, ectomycorrhiza versus control root. **(Separate excel file)**

**Table S5** *L. bicolor* genes specifically differentially expressed at only one of the time-points during the interaction (p-adj $< 0.01$; $\log_2$ fold change $> 0.5$) **(Separate excel file)**

**Table S6** *Populus* genes specifically differentially expressed at only one of the time-points during the interaction (p-adj $< 0.01$; $\log_2$ fold change $> 0.5$). **(Separate excel file)**

**Table S7** Differentially expressed (p-adj $< 0.01$; $\log_2$ fold change $> 0.5$) *L. bicolor* CAZYmes in our study compared to literature studies by Veneault-Fourrey *et al*. 2014 and Plett *et al*. 2015 (**Separate excel file**)

**Table S8** Differentially expressed (p-adj $< 0.01$; $\log_2$ fold change $> 0.5$) *P. tremula x P. tremuloides* CAZYmes in our study (**Separate excel file**)

**Methods S1** Bioinformatics

**Methods S2:** PME activity assay in free living mycelium

**Methods S3** Immunofluorescence localization of pectin antibodies in poplar roots

**Method S4** Cloning, *L. bicolor* transformation and fungal selection

**Method S5** Characterization of T-DNA insertion number in transgenic *L. bicolor* lines by ddPCR

**Method S6** Characterization of T-DNA insertion sites in transgenic *L. bicolor* by Plasmid rescue and TAIL-PCR

**Methods S7** Characterization of T-DNA insertion number in transgenic L. bicolor lines by ddPCR

**Fig. S1** Schematic diagram of vector constructs used in this study for altering *L. bicolor* PME expression.

*LbPME1* RNAi trigger expression cassette in pSILBAɣ

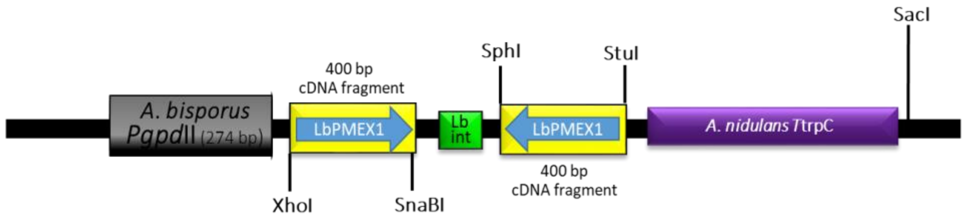


*LbPME* double RNAi trigger expression cassette in pSILBAɣ

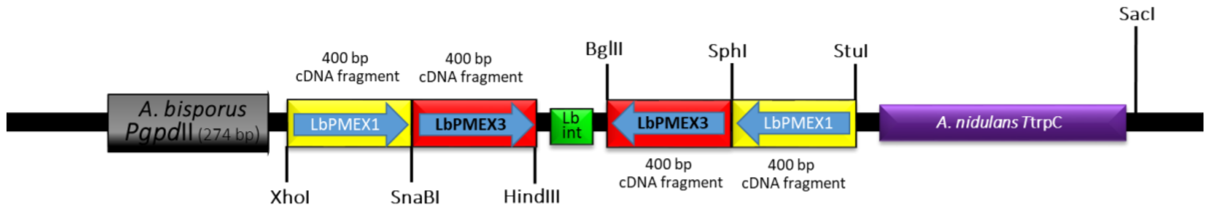


*LbPME1* over-expressor construct

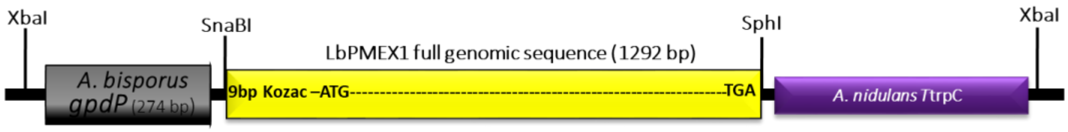

**Fig. S2** qPCR screening of *L. bicolor* RNAi and over-expressor (OE) transformant lines for *PME* expression in free-living mycelium relative to transcript levels in wild-type *L. bicolor*. For each construct, 12 to 14 transformed fungal lines were selected for qPCR screening. *LbPME2_4* indicates expression of *LbPME2*, *3* and *4* that are too close in sequence to distinguish them with qPCR. Lines marked * were selected for further characterization in this study.

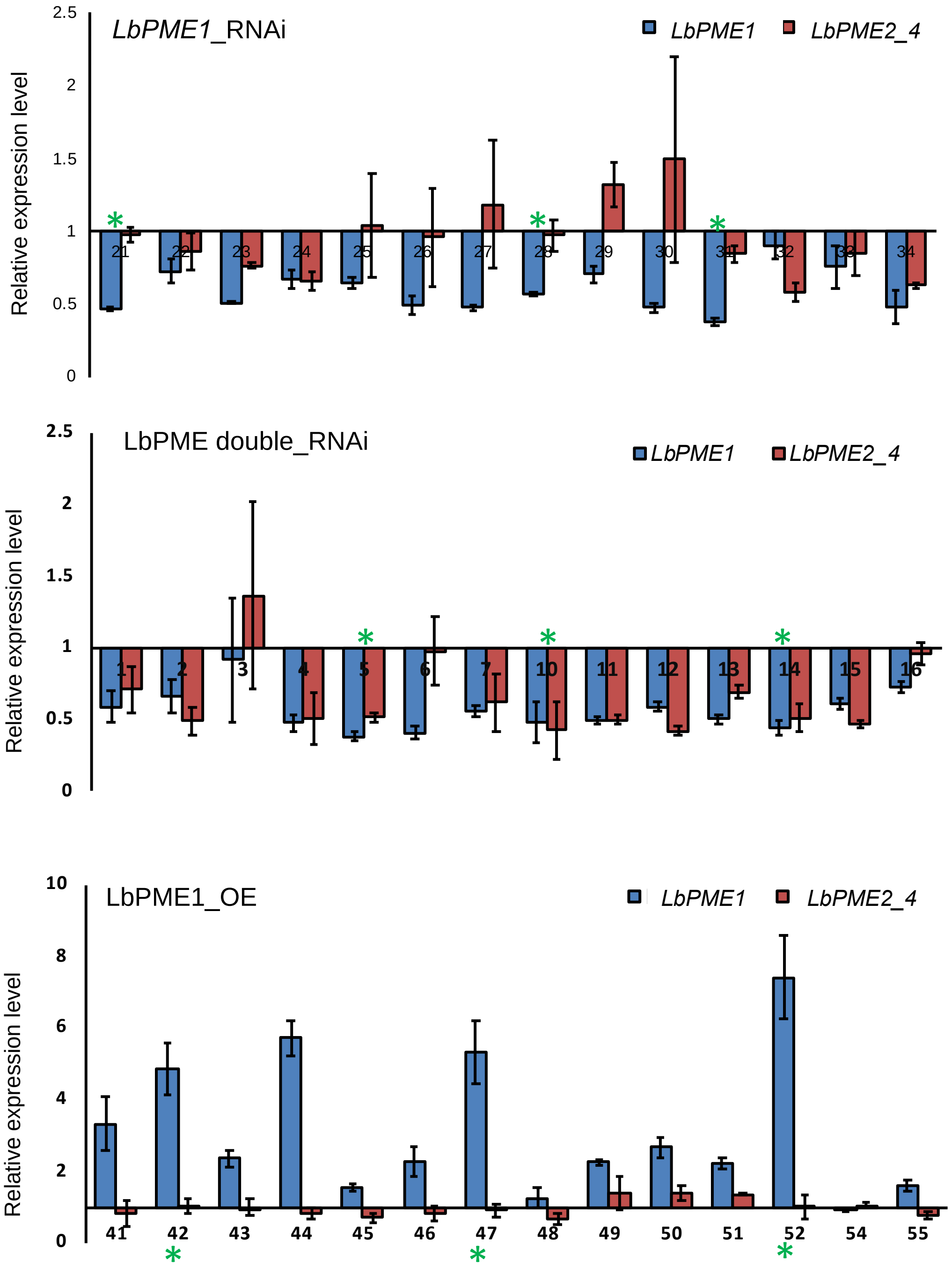

**Fig. S3** Gradual development of *L. bicolor* colonization while interacting with *P. tremula* x *P. tremuloides* lateral roots at various days after contact (DAC) under the *in vitro* sandwich culture system. The fungal cell-wall was stained with WGA-AF488 (green) and the plant cell-wall was stained with Pontamine fast scarlet 4BS (magenta). Scale bar = 20 $\mu$m

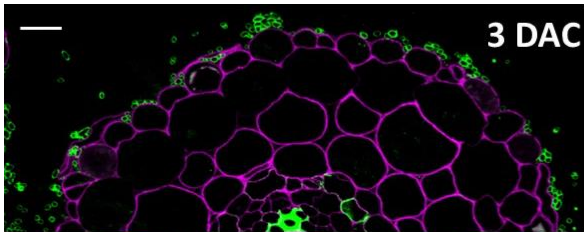


Early colonization phase of *L. bicolor* with *P. tremula x P. tremuloides*. A few hyphae (green) are starting to adhere to the root epidermis cells (magenta).

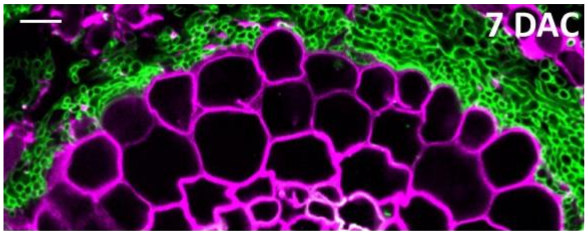


Mantle formation by *L. bicolor* with *P. tremula x P. tremuloides*. The Hartig Net is not formed yet.

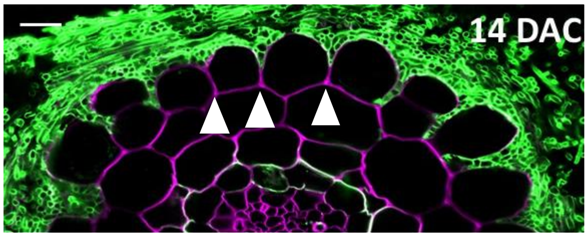


The Hartig Net is forming as seen by *L. bicolor* hyphae being present in between of epidermal cells of *P. tremula x P. tremuloides* roots. The penetration is limited to the epidermal layer and for some epidermal cells cell-separation is not yet complete (arrowheads).

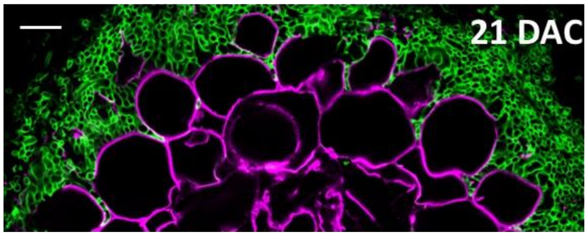


Mature ectomycorrhiza with a clearly visible Hartig Net. *L. bicolor* hyphae penetrate the entire epidermal layer of *P. tremula x P. tremuloides* and cell separation in the epidermal layer is complete.

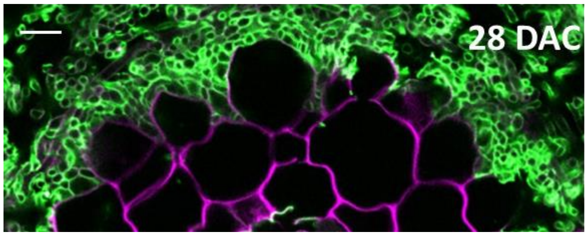


Late stage of ectomycorrhizal development. The number of fungal hyphae between root cells may increase. In *in vitro* culture conditions of *L. bicolor* and *P. tremula x P. tremuloides* root penetration usually stays limited to the epidermis layer and does not continue deeper into the cortex.

**Fig. S4** Venn diagrams of differentially expressed (DE) genes (p-adj < 0.01; $log_2$ fold change > 0.5) in mycelium in ECM compared to free-living mycelium (a) down and (b) up, as well as DEs in colonized *P. tremula* x *P. tremuloides* roots compared to roots without fungus (c) down and (d) up, at various time-points of development.

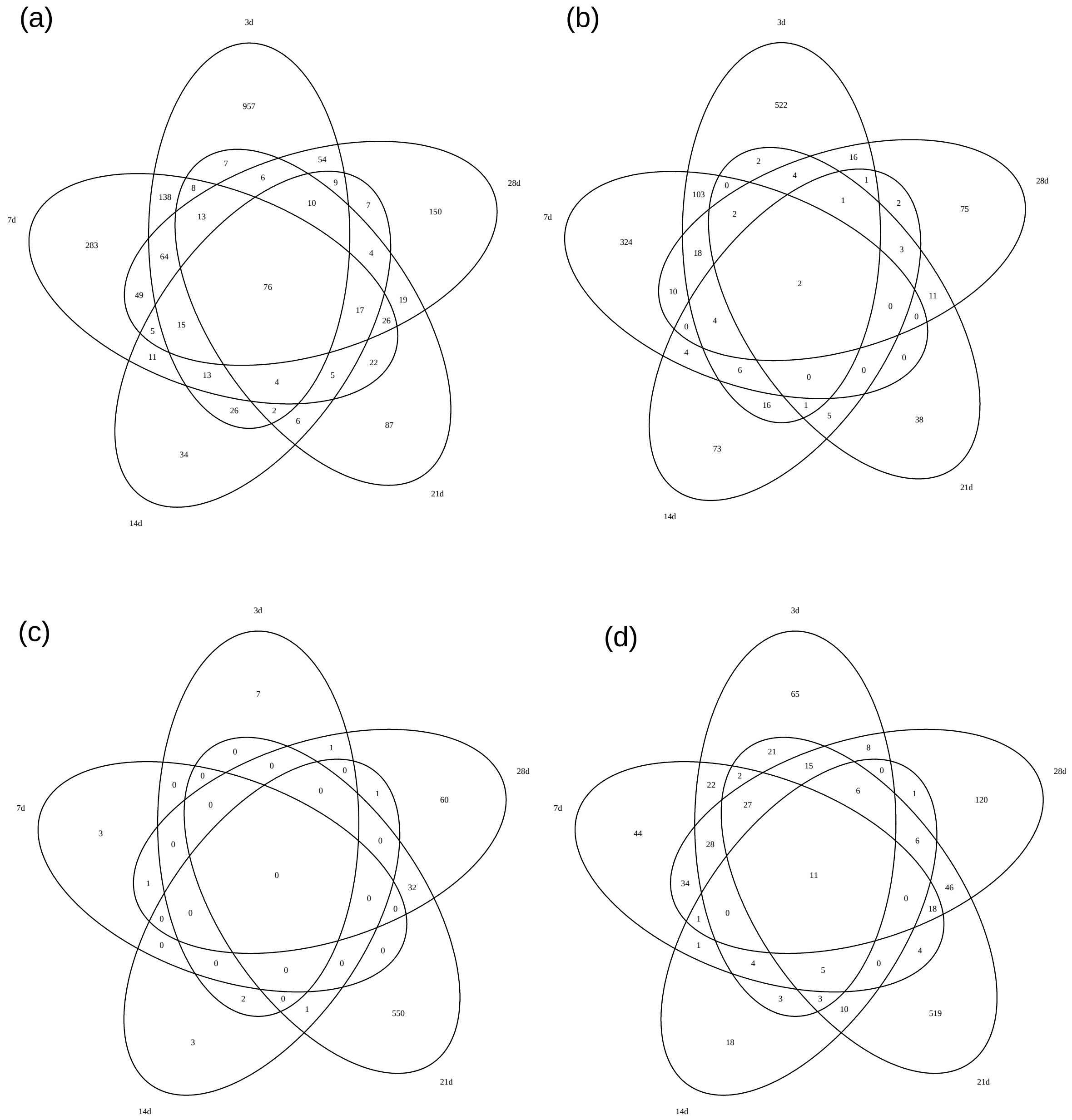

**Fig. S5** Proportion of differentially expressed genes and unchanged genes associated with specific polysaccharide degrading enzymes in *P. tremula* x *P. tremuloides* (a) and *L. bicolor* (b).

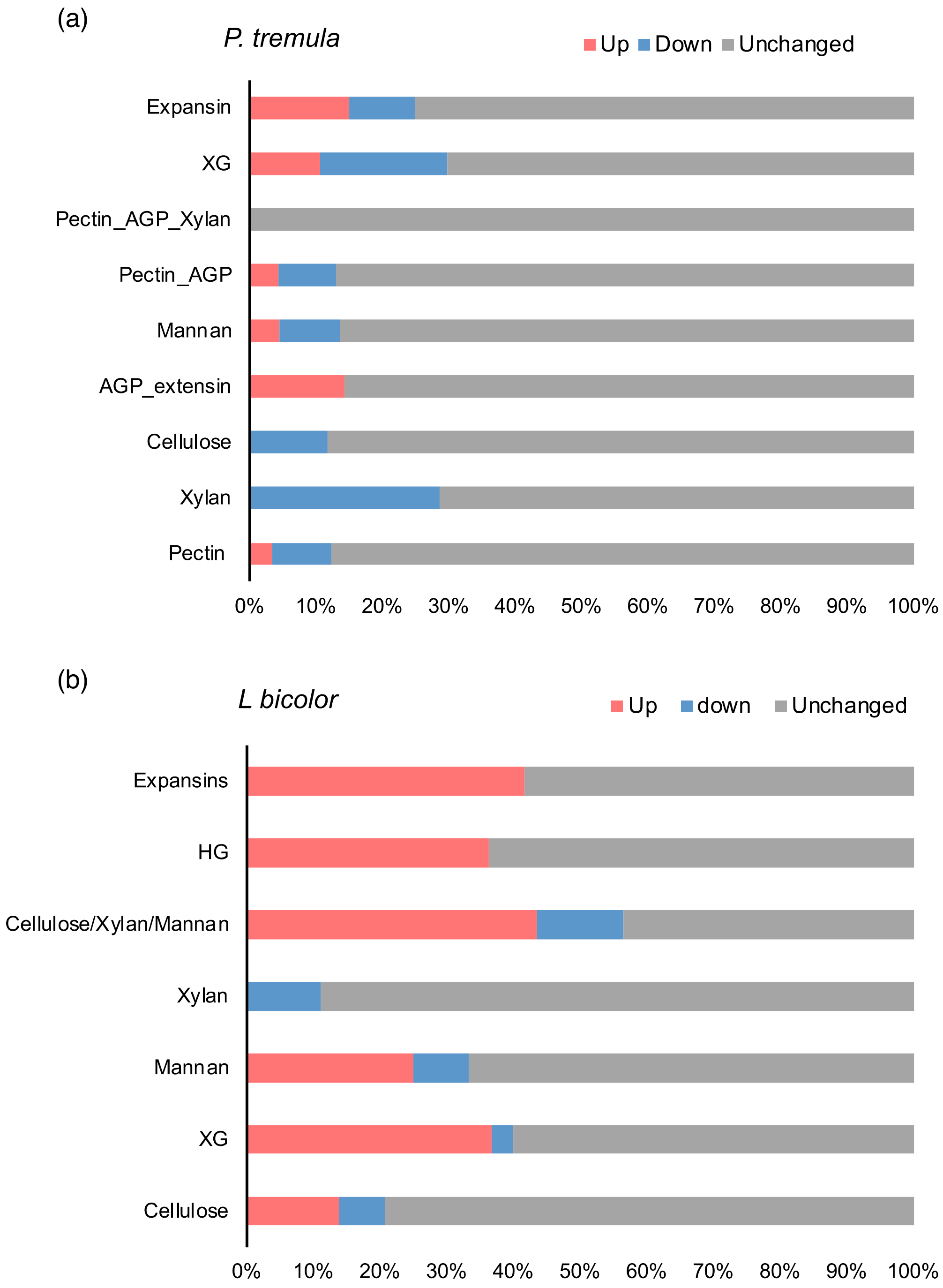

**Fig. S6** Time-course RNA-Seq expression profile of the potential candidates of HG synthesis and modifying enzymes (HGMEs) during *L. bicolor / P. tremula* x *P. tremuloides* interaction. Heat map represents $\log_2$ fold change ratios of differentially regulated genes ($\log_2$ fold change 0.5, padj < 0.05) in colonized roots versus control roots without fungus harvested at the respective similar timepoint. $\text{Log}_2$ Fold change values are indicated. PME = Pectin methylesterase, PMEI = Pectin methylesterase inhibitor, PAE = Pectin acetylesterase , PL = Pectate lyase, PG = Polygalacturonase, PGI = Polygalacturonase inhibitor, GT = Glycosyl transferase. Note that PMEI, PAE, PGI, and PL gene families are absent in *L. bicolor*. Figure on right shows a diagrammatic representation of homogalacturonan and the cleavage and addition (GT8) sites of potential HGMEs.

| | 3d | 7d | 14d | 21d | 28d |
|---|---|---|---|---|---|
| **LbPMEs** | | | | | |
| Lb 245379 | 3.9 | 1.6 | 2.2 | 1.7 | 1.9 |
| **LbPGs** | | | | | |
| Lb613299 | 2.6 | ns | 1.2 | 1.6 | 1.3 |
| Lb388191 | 2.5 | ns | ns | ns | ns |
| Lb612983 | 3.1 | ns | 1.9 | ns | 1.8 |
| **PtPMEs** | | | | | |
| Potra000192g00806 | ns | ns | ns | -6.5 | -6.3 |
| Potra172988g27617 | ns | ns | ns | -4.8 | ns |
| Potra000942g07708 | ns | ns | ns | -3.8 | ns |
| Potra000375g01594 | ns | ns | ns | -3.6 | ns |
| Potra001161g10080 | ns | ns | ns | -2.8 | ns |
| Potra001085g09413 | ns | ns | ns | -2.8 | ns |
| Potra003529g21908 | ns | ns | ns | -2.5 | ns |
| Potra001147g09998 | 1.2 | ns | 0.6 | ns | ns |
| Potra170651g27533 | ns | 3.0 | 3.2 | ns | ns |
| Potra000813g06484 | ns | 2.1 | ns | ns | ns |
| **PtPMEIs** | | | | | |
| Potra001916g15199 | ns | ns | ns | -5.1 | -3.4 |
| Potra000375g01597 | ns | ns | ns | -4.4 | ns |
| Potra009294g26327 | ns | ns | ns | 1.2 | ns |
| Potra001270g10974 | ns | ns | ns | 1.4 | ns |
| Potra154746g26750 | ns | ns | ns | ns | -1.8 |
| **PtPAEs** | | | | | |
| Potra000521g03520 | ns | ns | ns | -4.6 | ns |
| Potra000499g03218 | ns | ns | ns | -3.3 | ns |
| Potra002589g19469 | ns | ns | ns | -2.4 | ns |
| Potra000991g08212 | ns | ns | ns | -1.0 | ns |
| Potra000521g03518 | ns | 2.2 | ns | ns | ns |
| **PtPGs** | | | | | |
| Potra000948g07768 | ns | 1.5 | ns | ns | ns |
| **PtPGIs** | | | | | |
| Potra002267g17383 | ns | ns | 2.4 | 0.6 | ns |
| Potra000979g08101 | 2.2 | ns | 2.4 | 1.7 | ns |
| Potra001285g11091 | 3.8 | ns | ns | 2.7 | ns |
| **PtPL** | | | | | |
| Potra003212g21010 | ns | ns | ns | -4.1 | ns |
| Potra004445g24927 | ns | ns | ns | -2.7 | ns |
| Potra001572g13018 | ns | ns | ns | -2.4 | ns |
| Potra002262g17355 | ns | ns | ns | -1.9 | ns |
| Potra001311g11269 | ns | 1.8 | ns | ns | ns |
| **PtGT8** | | | | | |
| Potra001085g09395 | ns | ns | ns | -2.0 | ns |
| Potra001269g10964 | ns | ns | ns | -1.5 | ns |
| Potra002029g15886 | ns | ns | ns | -3.1 | ns |
| Potra002583g19425 | ns | ns | ns | -1.2 | ns |
| Potra175700g27820 | ns | 1.9 | ns | 1.3 | 2.6 |

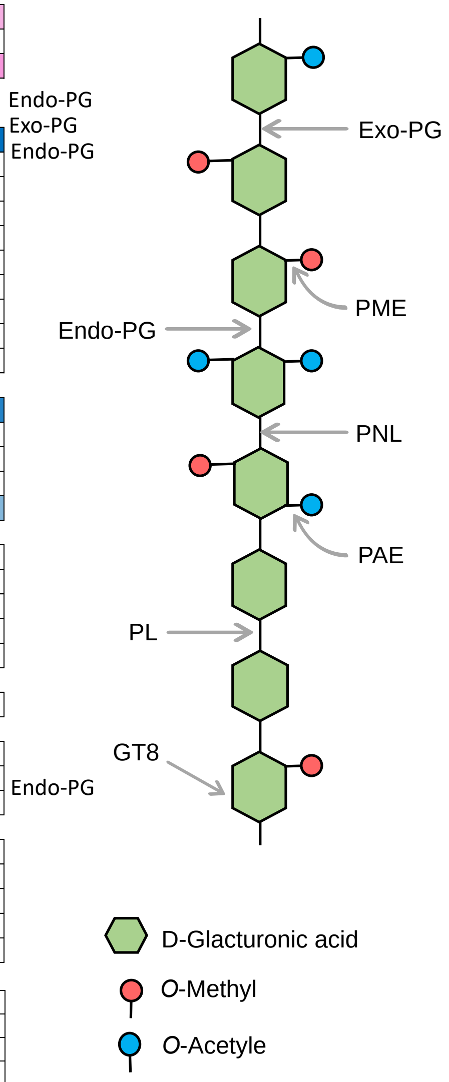

**Fig. S7** Comparison of gene expression values obtained by RT-qPCR and RNA-seq. $Log_2$ Fold changes in colonized roots were calculated for two differentially expressed genes from plant (*PttMYB134*) and fungus (*LbPME1*) compared to control roots and FLM, respectively (a). For qPCR, expression value was normalized using three stable housekeeping genes. A high correlation ($R^2$ > 0.99) was observed between the results obtained using the two techniques (b).

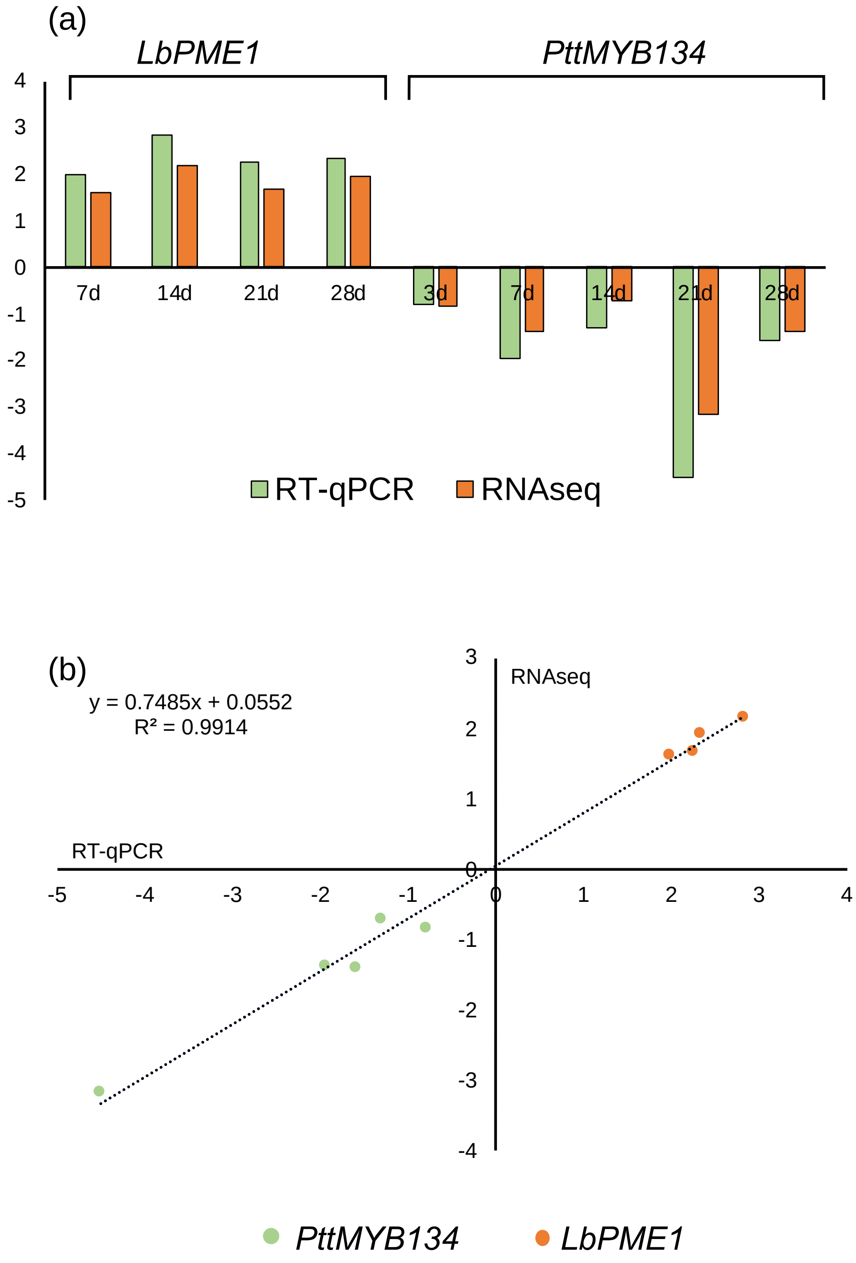

**Fig. S8** Time-course RNA-Seq expression profile of four *L. bicolor* PMEs in colonized roots (ECM) and free-living mycelia (FLM).

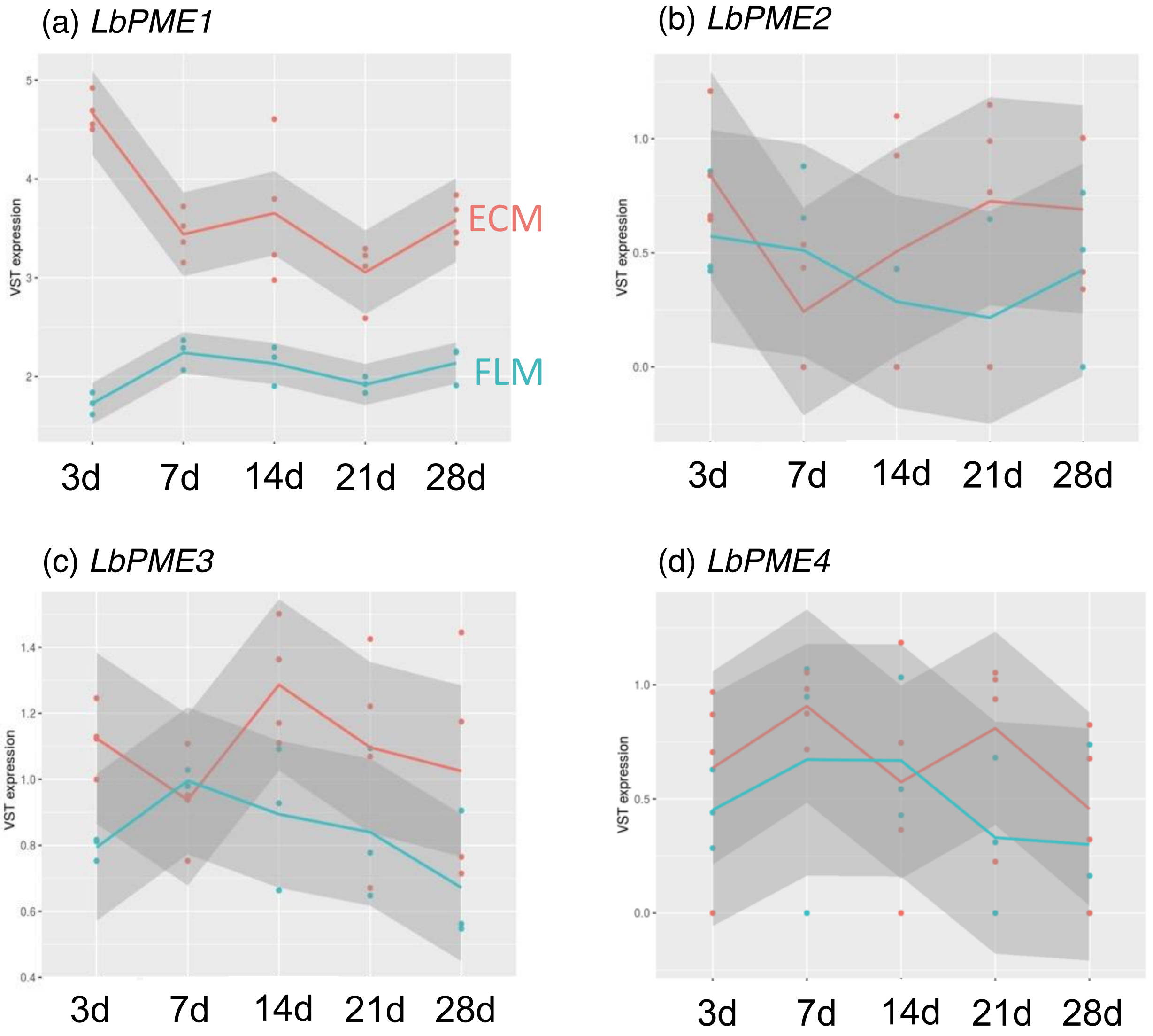

**Fig S9:** Differential expression of *P. tremula x P. tremuloides* HG biosynthesis genes during interaction with *L. bicolor* based on RNASeq data from our study. (a) Differentially expressed genes of the GT8 (GAUT and GATL) family. Heat map represents $\log_2$ fold change ratios of differentially regulated genes ($\log_2$ fold change 0.5, padj < 0.05) in colonized roots versus control roots without fungus harvested at the respective similar time-point. $\text{Log}_2$ Fold change values are indicated. (b) Identified pectin methyltransferases in *Populus*, based on homology. None of these were differentially expressed.

(a)

| GT8 | 3d | 7d | 14d | 21d | 28d | Poplar homolog | Gene ID | Gene annotation |
|---|---|---|---|---|---|---|---|---|
| Potra001085g09395 | ns | ns | ns | -2.0 | ns | Potri.002G200200 | PtGATL7-A | Probable galacturonosyltransferase-like |
| Potra001269g10964 | ns | ns | ns | -1.5 | ns | Potri.014G073800 | GAUT1 | 4-alpha-galacturonosyltransferase |
| Potra002029g15886 | ns | ns | ns | -3.1 | ns | Potri.014G029900 | GT8 | plant glycogenin-like starch initiation protein 3 |
| Potra002583g19425 | ns | ns | ns | -1.2 | ns | Potri.014G125000 | PtGATL7-B | Probable galacturonosyltransferase-like |
| Potra175700g27820 | ns | 1.9 | ns | 1.3 | 2.6 | Potri.008G192600 | PtGATL8-A | Probable galacturonosyltransferase-like |

(b)

| Gene | Poplar homolog | ATG homolog | Gene annotation |
|---|---|---|---|
| Potra001251g10774 | Potri.001G288200.1 | AT3G49720.1 | Uncharacterized protein At3g49720 |
| Potra002036g15919 | Potri.009G082400.1 | AT1G13860.4 | Probable methyltransferase PMT4 |
| Potra002518g19016 | Potri.008G094800.1 | AT2G03480.1 | Probable methyltransferase PMT5 |
| Potra176014g27856 | Potri.010G159400.1 | AT4G00740.1 | Probable methyltransferase PMT13 |

**Fig. S10** Comparison of differentially expressed Lb CAZymes in our study to those in Veneault-Fourrey *et al*. 2014 (a) and Plett *et al*. 2015 (b) during interaction of *Populus* with *L. bicolor* in a soil system. 63 differentially expressed CAZymes (pVal < 0.05, $log_2$ fold change > 0.5) were compared to the published data. The clusters into which the respective publications sorted the expressed genes are listed. The percentage indicates the proportion that the gene number comprises within the respective cluster.

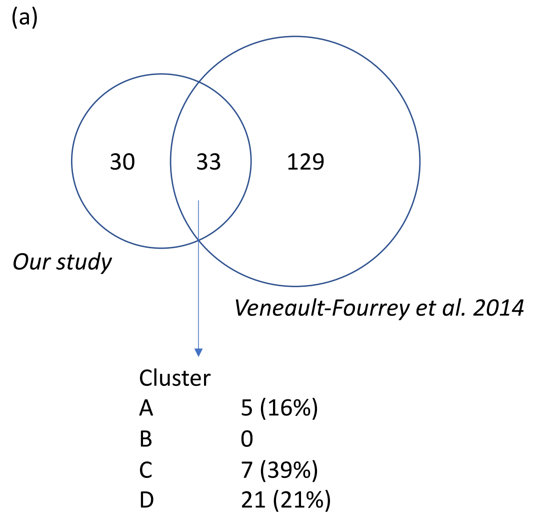


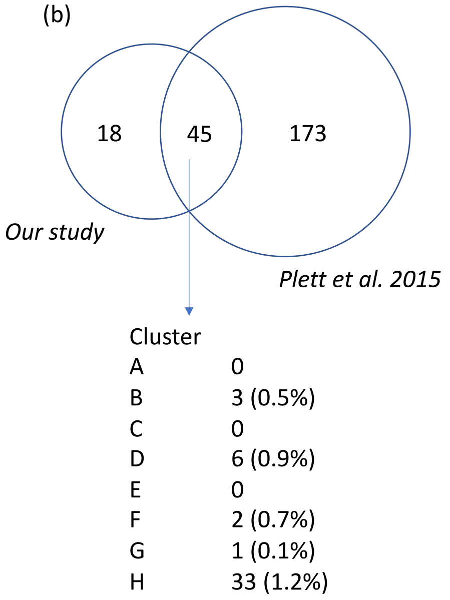

**Fig. S11** Immunolocalization with pectin antibodies LM19 and LM20 at various distances from the root tips. Representative visualization of antibody labeling at a specific distance from the tips of colonized roots (magenta) with *L. bicolor* (green) (a) and control *P. tremula* x *P. tremuloides* roots (b). LM19 and LM20 fluorescence intensity of Hartig Net wall for colonized roots and radially adjacent cell-wall in control roots (c). At least five root tips collected from five plants at 14 DAC were examined. Measurements were made at 10 positions per micrograph and averaged. Statistical analysis was performed using paired Student t-test. P value was < 0.05 for all positions.

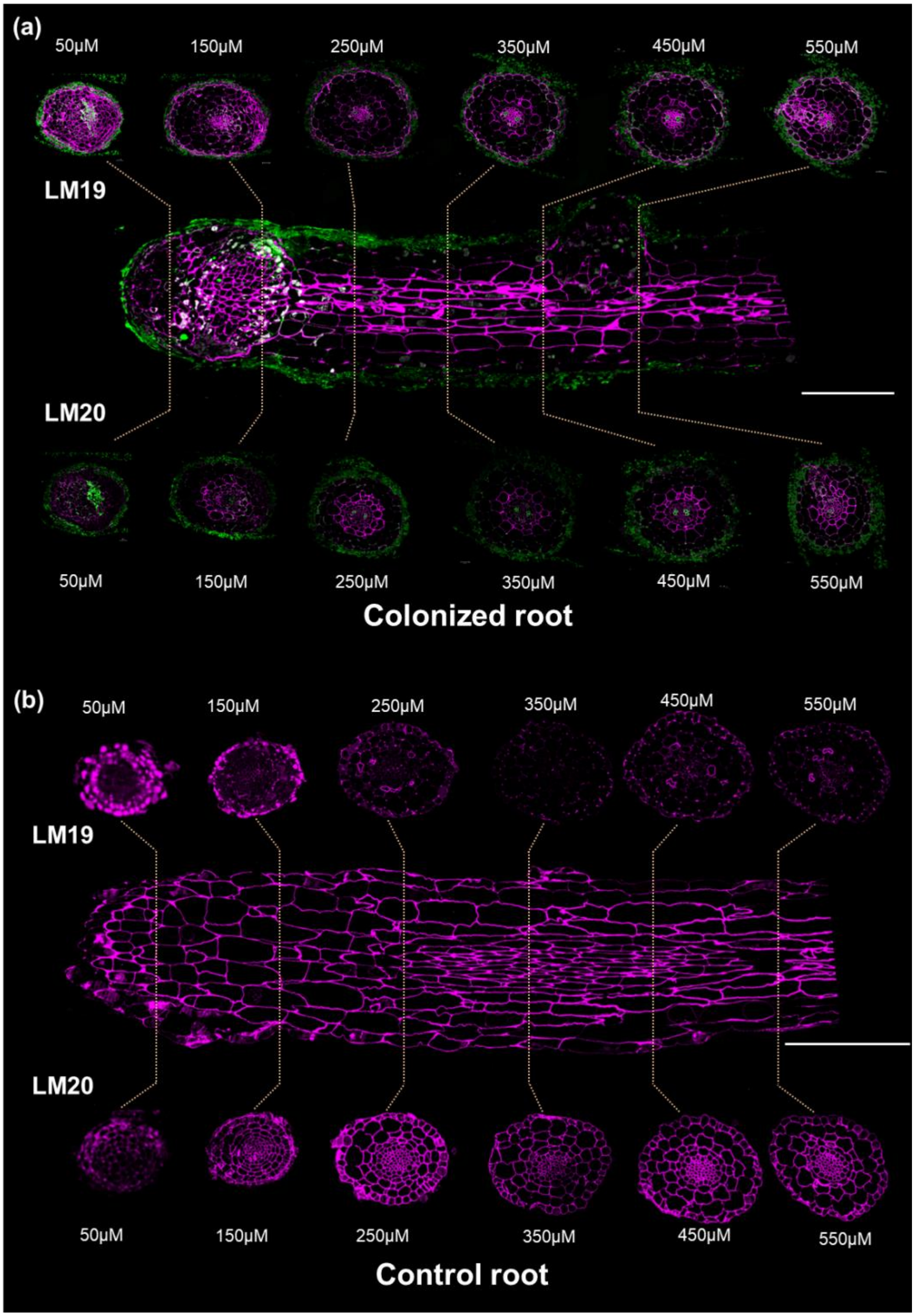

**Fig. S11** continued

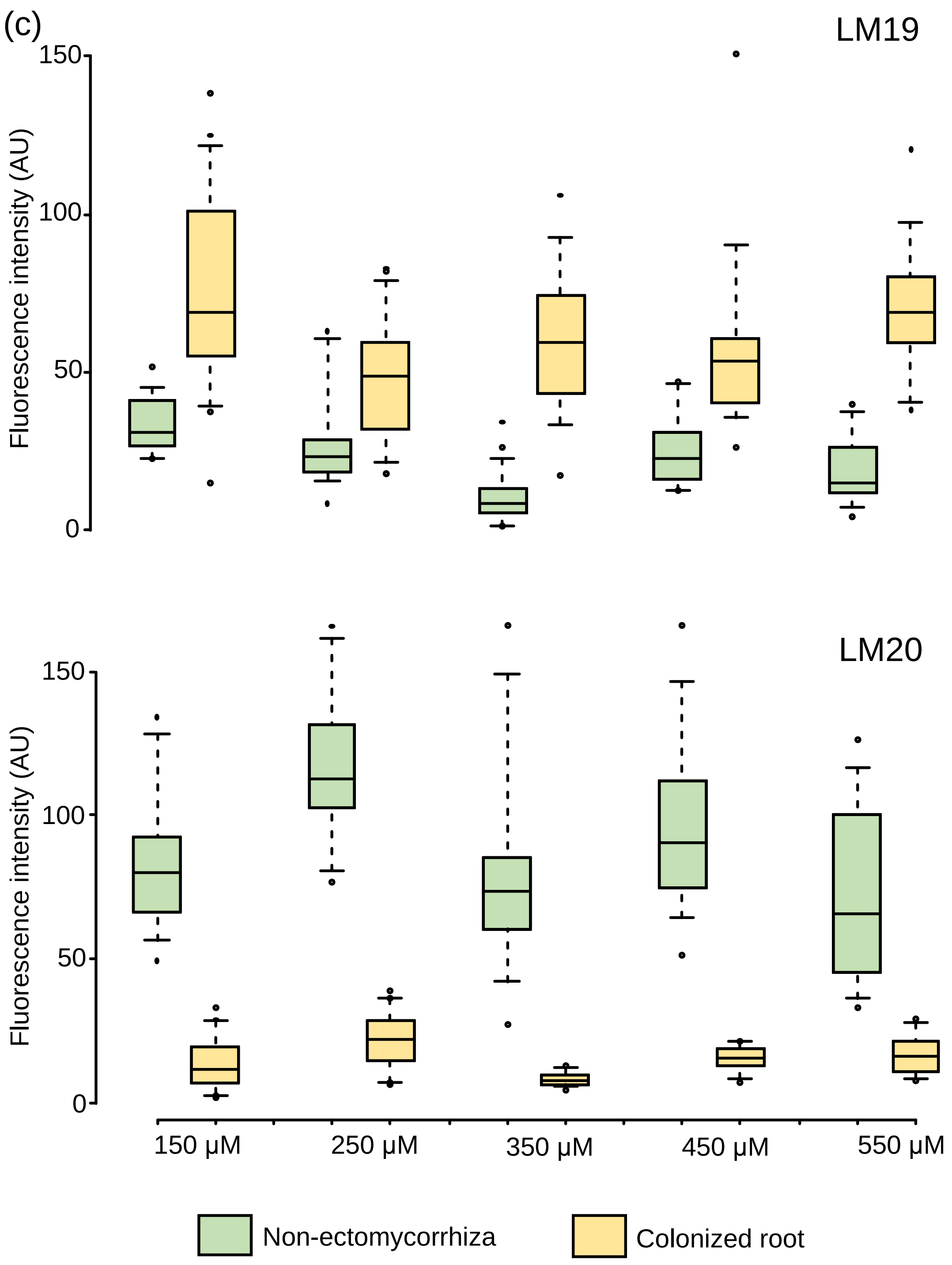

**Fig. S12** Transmission electron micrograph of immuno-gold labeling of homogalacturonan epitopes present in the epidermis cell-wall of ECM (a,b,e,f) and control root cross-sections (c,d,g,h). Samples were collected at 14 DAC and were embedded in LR White resin. The epitopes of low methylesterified and high methylesterified homogalacturonan were detected using LM19 (a-d) and LM20 antibody (e-h). Gold particles are arrowed. Quantitative data are available in Fig. S13.

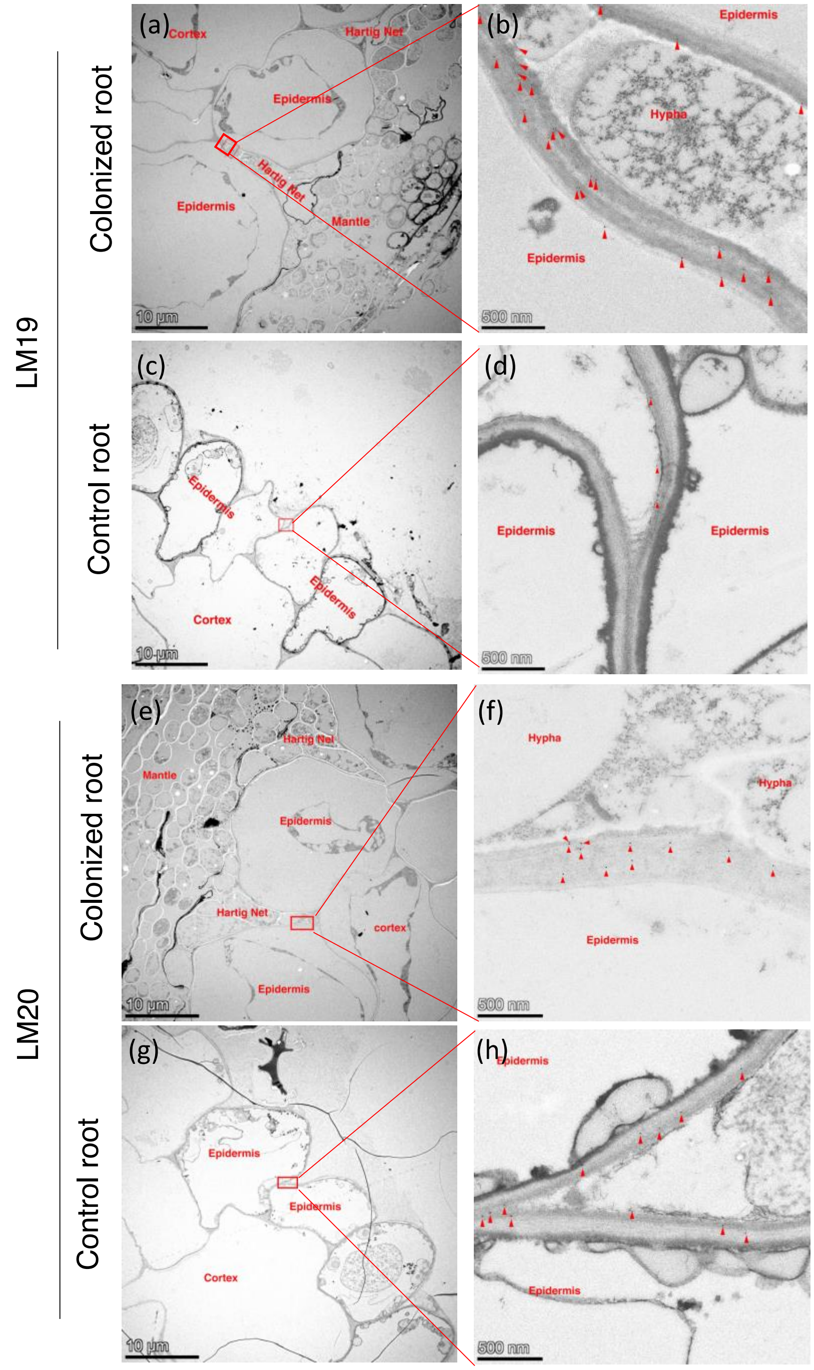

**Fig. S13** Quantification of gold particles in the Hartig Net plant cell-wall and corresponding cell-wall areas of control roots on TEM images (examples shown in Fig S12 b, d, f, h). For each treatment, 10-16 images at x22 000 magnification from three individual root section were analyzed per antibody. Statistical analysis was performed using paired Student t-test. **P value < 0.01.

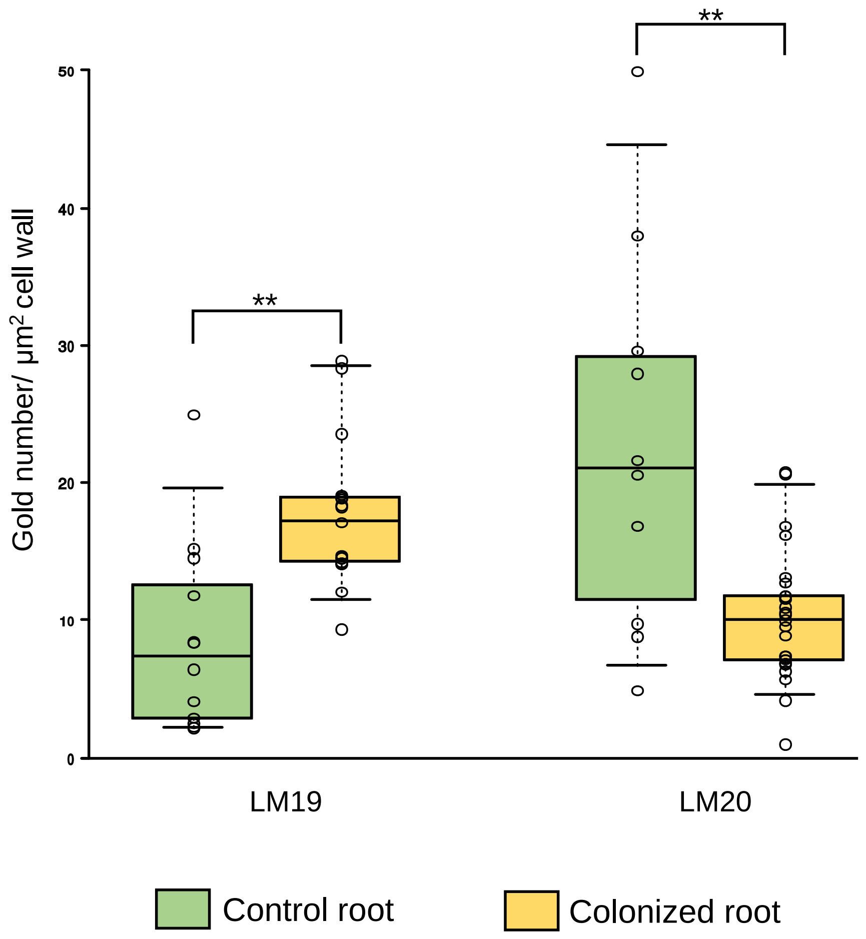

**Fig. S14** Multiple sequence alignment (a) and sequence identity (b) of *L. bicolor* PME proteins. Conserved amino acid residues are indicated by black shading whereas similar residues are represented in grey shading using a threshold for shading of 60% similarity (a). All four proteins consist of signal peptides (red bracket) with no transmembrane domain and a PME catalytic domain (Pfam01095, blue bracket) suggesting that they belong to Group 1 (or type 2) class of PMEs (Pelloux *et al.*, 2007; Sénéchal *et al.*, 2015).

**(a)**

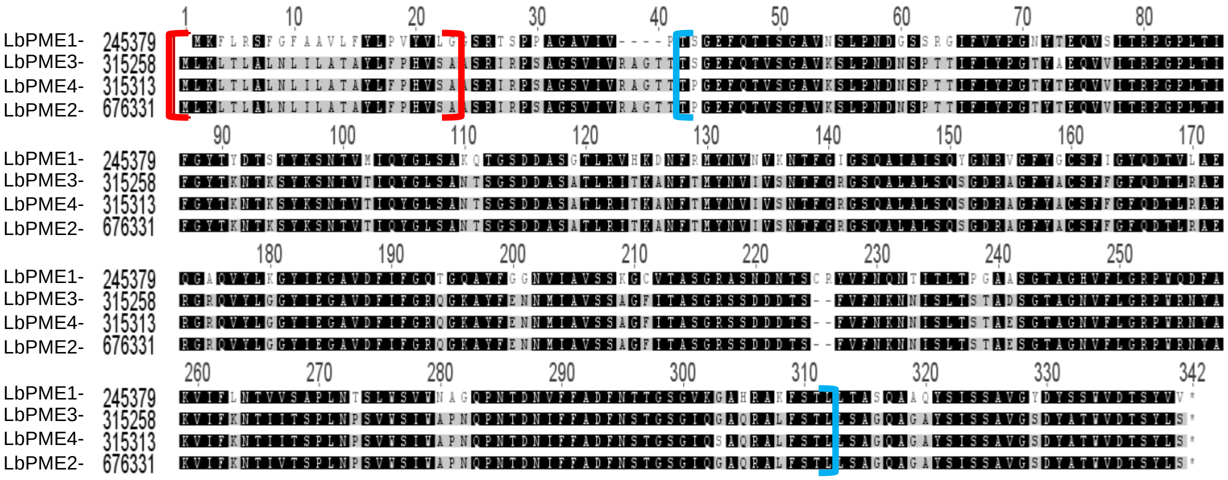


**(b)**

| | PME1 | PME3 | PME4 | PME2 |
|---|---|---|---|---|
| **PME1** | | 65% | 65% | 66% |
| **PME3** | | | 99% | 99% |
| **PME4** | | | | 99% |
| **PME2** | | | | |

**Fig S15** Relative normalized expression of *LbPGs* in free-living mycelium of the PME transgenic lines by qPCR (n = 3 biological replicates). EV7 and EV9 are empty vector controls. Statistical analysis was performed using a Student T-Test compared to wild-type fungus (WT). *P < 0.05. Error bars show standard deviation of biological replicates. The dotted line shows the average expression of the genes in WT.

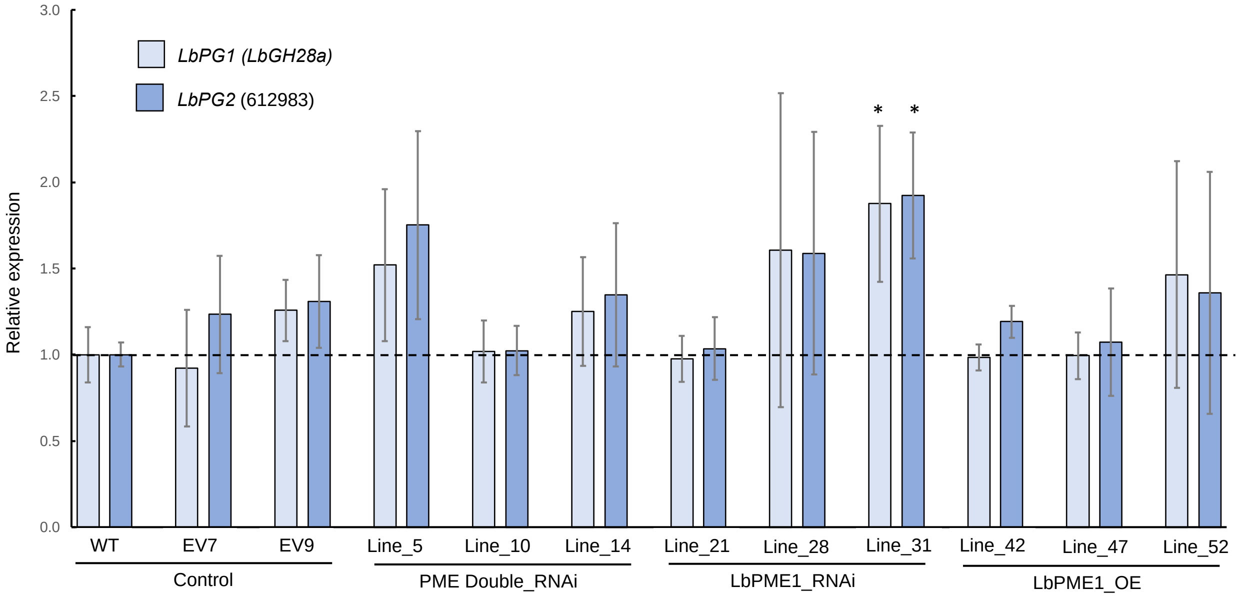

**Fig. S16** Correlation of *PME1* transcript level and PME enzymatic activity level in the free-living mycelia of the transgenic fungal lines. Transcript level and PME activity level are relative to the respective wild-type free-living mycelium control.

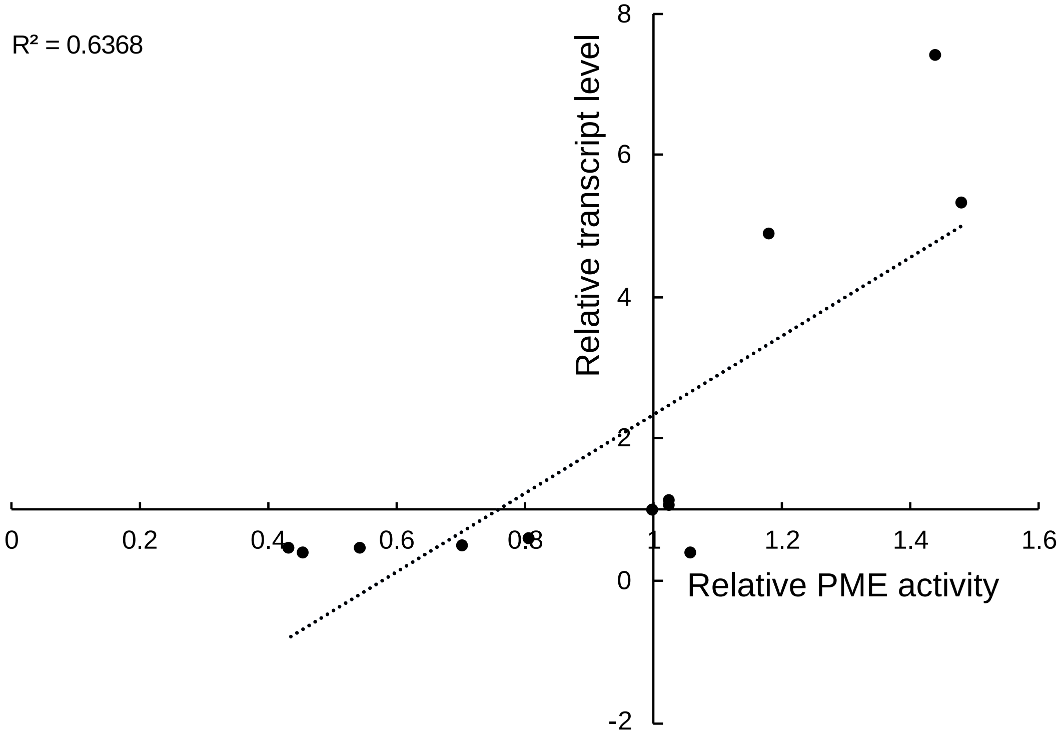

**Fig. S17** Examples of control roots (a, b) and swollen root tips with attachment of fungal mycelia (c, d) on colonized roots, which are considered as ectomycorrhizal root tips. Blue dots result from experimental practice of ECM counting and are to be ignored. (b) is a magnification of (a), and (d) is a magnification of (c).

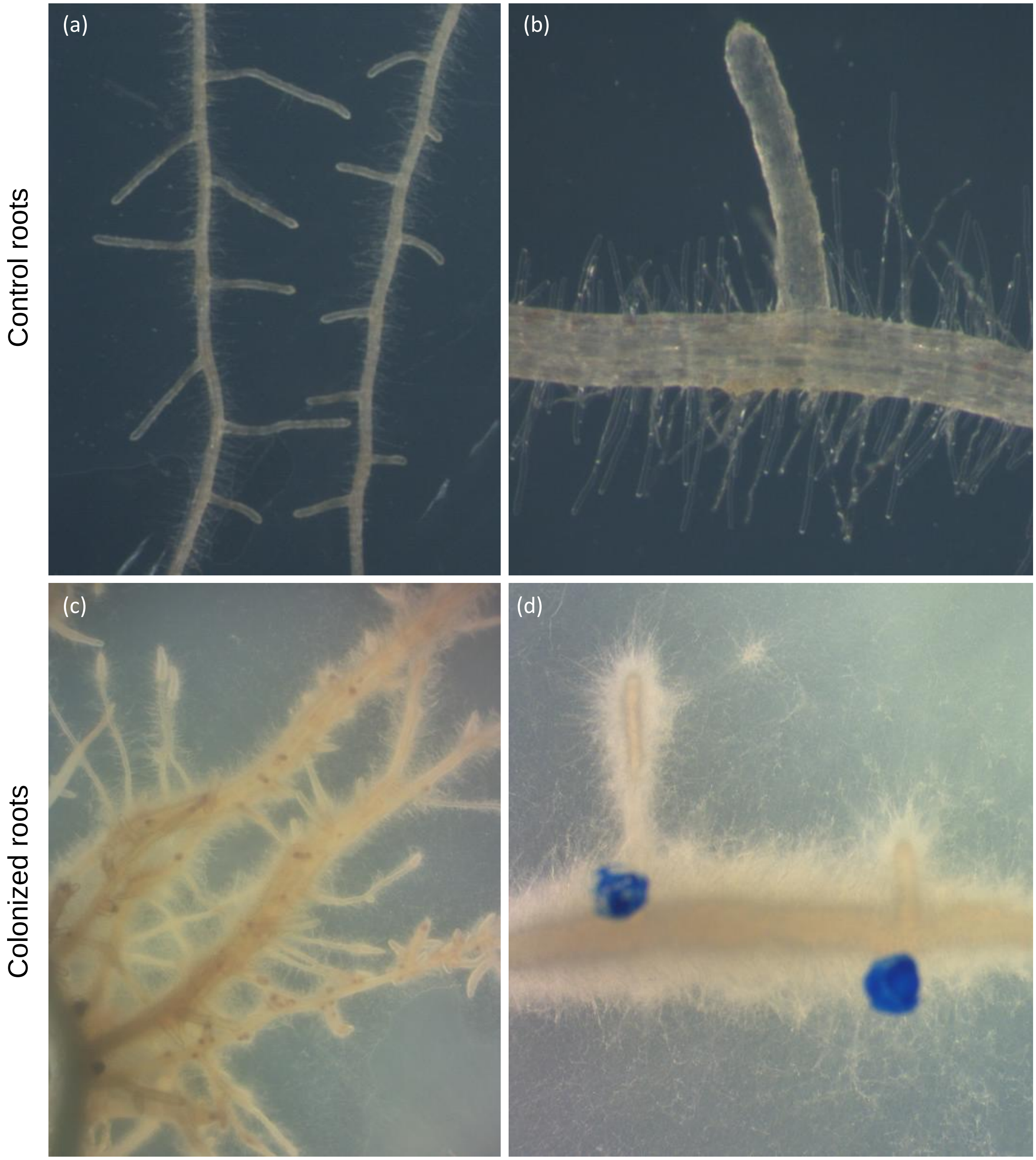

**Fig. S18** Representative micrographs of sections through ectomycorrhizal root tips of *P. tremula x P. tremuloides* in contact (14 days) with *L. bicolor* wild-type (a), empty vector line 7 (b), PME1 RNAi Line 21 (c), PME double RNAi Line 10 (d) and PME1 overexpressor Line 47 (e). WGA-Alexa 488 stain for fungus (green), and Pontamine Fast Scarlet stain for plant cells (magenta). Scale bar = 20 $\mu$m. (f) LM19 (CY5) fluorescence intensity in radial epidermal walls. Large dots represent the mean of individual lines (same lines as e.g. Fig S15). Small dots represent individual fluorescence intensity measurements (n=15-20 with 3-4 ECM per line and 5-6 measurements per micrograph). Horizontal lines show mean and standard deviation of the entire observation per category. P-values were calculated based on Student-T-Test against controls (comprising of empty vector line 7 and 9 and wild-type). The plot was created using SuperPlotsOfData Shiny app (Goedhart, 2021).

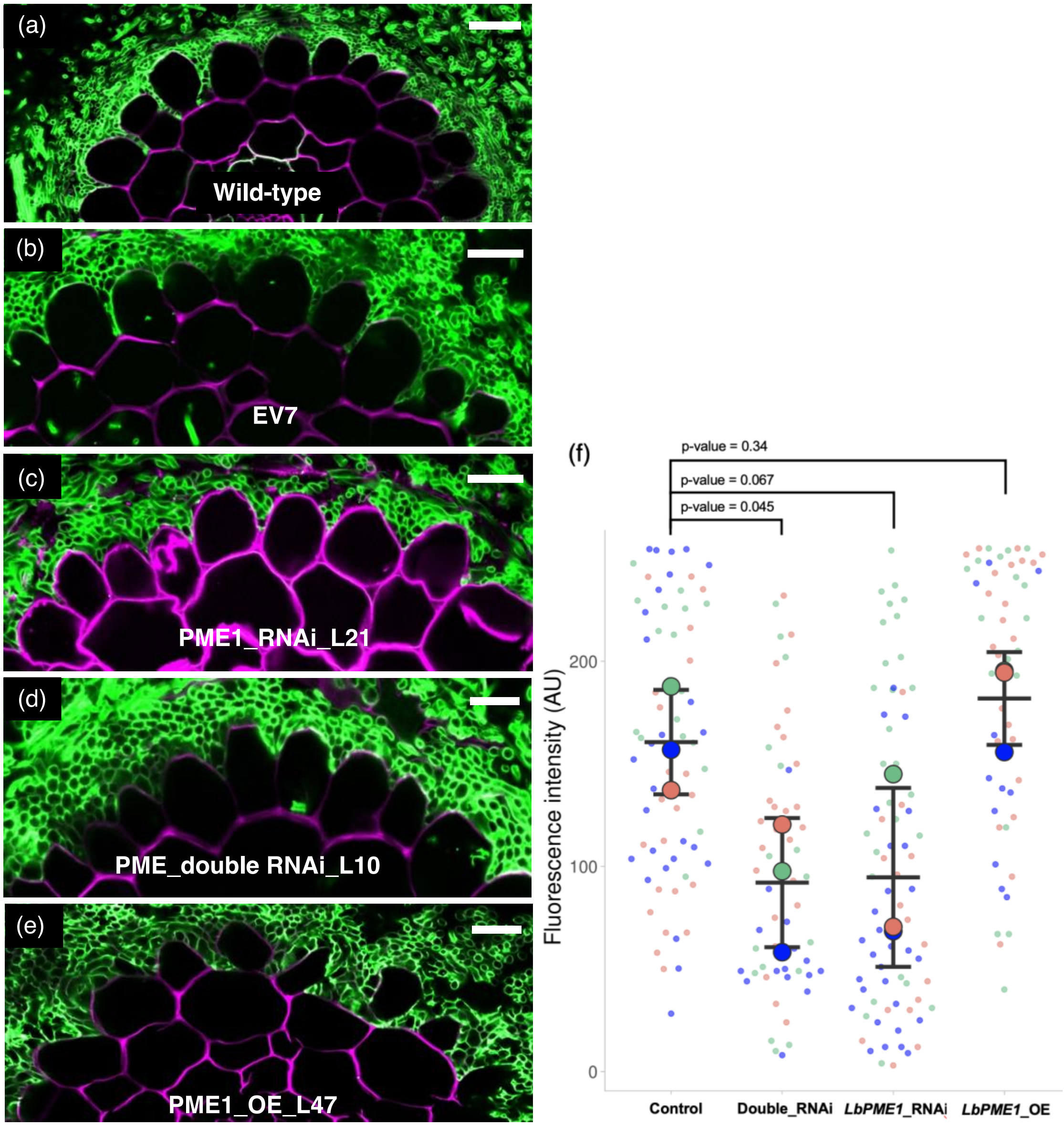

**Methods S1** Bioinformatics

The quality of the raw sequence data was assessed using FastQC (http://www.bioinformatics.babraham.ac.uk/projects/fastqc/). Residual ribosomal RNA (rRNA) contamination was assessed and filtered using SortMeRNA (v2.1;(Kopylova *et al.*, 2012); settings --log --paired_in --fastx--sam --num_alignments 1) using the rRNA sequences provided with SortMeRNA (rfam-5s-database-id98.fasta, rfam-5.8s-database-id98.fasta, silva-arc-16s-database-id95.fasta, silva-bac-16s-database-id85.fasta, silva-euk-18s-database-id95.fasta, silva-arc-23s-database-id98.fasta, silva-bac-23s-database-id98.fasta and silva-euk-28s-database-id98.fasta). Data were then filtered to remove adapters and trimmed for quality using Trimmomatic (v0.36; (Bolger et al., 2014); settings TruSeq3-PE-2.fa:2:30:10 LEADING:3 SLIDINGWINDOW:5:20 MINLEN:50). After both filtering steps, FastQC was run again to ensure that there were no introduced technical artefacts. Filtered reads were pseudo-aligned to v1.1 of the *P. tremula* transcripts (closest reliable genome to *P. tremula* x *P. tremuloides*, retrieved from the PlantGenIE resource; (Sundell *et al.*, 2015) using Salmon (v0.14.1; (Patro *et al.*, 2017); non default settings: --gcBias --seqBias). Statistical analysis of single-gene differential expression between conditions was performed in R (v3.6.2; R Core Team 2019) using the Bioconductor (v3.9; (Gentleman *et al.*, 2004). DESeq2 package (v1.26. 0; (Love *et al.*, 2014). FDR adjusted p-values were used to assess significance; a common threshold of 1% was used throughout. For the data quality assessment (QA) and visualisation, the read counts were normalized using a variance stabilizing transformation as implemented in DESeq2. The biological relevance of the data - e.g. biological replicates similarity - was assessed by Principal Component Analysis (PCA) and other visualizations (e.g. heatmaps), using custom R scripts, available from https://github.com/nicolasDelhomme/laccariaBicolorEcmDev.

**Methods S2** PME activity assay in free-living mycelium

Approximately 100 mg of free-living mycelia were grown on modified P20 media containing *P. tremula* x *P. tremuloides* root cell-wall materials as the carbon source (obtained from roots grown on cellophane membranes on half-strength MS media, frozen and freeze-dried, incorporated at 2%, w/w dry basis into the media for this experiment). Fourteen-day old FLM were harvested and flash-frozen in liquid nitrogen. Total protein was extracted from the finely ground samples by suspending in extraction buffer containing 100 mM Tris-HCl (pH 7.0), 500 mM NaCl and 1x protease inhibitor cocktail (abmGood, Cat. number: G135) at 4 °C for one hour with repeated vortexing (20 s every 10 minutes). Extracts were centrifuged (14,000g, 30 min at 4 °C) and the total protein (supernatant) was collected. Then 50 µL of total protein (50 ng/uL) from each sample were incubated with 75 µL of MBTH (N-methylbenzothiazolinone-2-hydrazone) solution (20 µL 100 mMTris-HCl, pH 7.5 containing 0.5% methylesterified pectin, 2 µL alcohol oxidase (0.02 U/µL), 8 µL MBTH (3mg/mL in water)) for 10 minutes at 40 °C. For colour development, the samples were then incubated with 100 µL solution containing ferric ammonium sulphate (5 mg/mL in water) and sulfamic acid (5 mg/mL in water). After 15 minutes at room temperature, every sample was diluted with 150 µL of water before measuring the absorbance at 620 nm using a microplate reader (Epoch, Biotek, USA). Four biological replicates and two technical replicates were analysed for each line using Fisher's LSD test. We performed two independent PME activity assays of FLM and obtained similar results.

**Methods S3** Immunofluorescence localization of pectin antibodies in poplar roots

Three hydrophobic circles were drawn using a PAP pen around the three separately placed slide-mounted sections. The sections were then rehydrated with Phosphate-buffered saline (PBS) and incubated with PBS containing 10µg/ml wheat germ agglutinin (WGA) conjugated with Alexa Fluor 488 Conjugate (ThermoFisher Scientific) for 30 minutes to label the fungal cell-wall. After washing with PBS, the slides were incubated in PBS containing 0.05 M glycine for 20 minutes and blocked with a blocking buffer containing 2% (w/v) bovine serum albumin (BSA) in PBS for 30 minutes. LM19 or LM20 primary antibody (diluted 1:20 in PBS) was added in two circles separately and only PBS was added in the third circle, which was used as the negative control. After incubating for two hours in a humid chamber at room temperature, the slides were washed three times with blocking buffer. All sections including the negative controls were incubated with Cy™5 AffiniPure Donkey Anti-Rat IgG (H+L) secondary antibody (Cat. Number 712-175-153, Jackson ImmunoResearch Europe Ltd.) for one hour at 1:100 dilution. Sections were washed three times with PBS and mounted in an antifade mounting media (CitiFluor™ AF1).

**Methods S4** Cloning, *L. bicolor* transformation and fungal selection

We amplified the full length CDS of *LbPME1*, *LbPME3* using pENTR™ Directional TOPO® Cloning Kit (Thermofisher Scientific) with primers given in Table **S1**. To construct the *LbPME1*

RNAi vector, a 400 bp fragment was amplified with two sets of primer pairs containing the restriction enzyme sites used in the subsequent cloning steps (Table **S1**). The two cDNA fragment arms were cloned into the pSILBAγ vector to complete the *LbPME1* RNAi construct, which results in the expression of an intron hairpin RNA (ihpRNA) (Kemppainen & Pardo, 2010) (Fig. S1). To silence all three *LbPMEs* simultaneously with a double-RNAi construct, a 400 bp cDNA fragment for RNAi was selected that is 99.25% identical between *LbPME3* and the two other *LbPMEs*. The two fragment arms were amplified from *LbPME3* full-length cDNA and cloned into pSILBAγ to construct the *LbPMEX3* RNAi vector. To complete the double trigger RNAi construct, the inverted sequence repeat was liberated from the *LbPMEX3* RNAi vector with SnaBI and SphI double digestion and cloned into equally SnaBI and SphI digested *LbPME1* RNAi vector (Fig S1). For the *Agrobacterium*-mediated transformation, the *LbPME1* and the *LbPME* double trigger RNAi vector were further cloned as full SacI linearized plasmids into the T-DNA of the binary vector pHg (Kemppainen & Pardo, 2010).

We also constructed *LbPME1* over-expressor lines. The *LbPME1* overexpression cassette was constructed in pSILBAγ vector under control of the *Agaricus bisporus GPD* gene promoter using the genomic sequence of *LbPME1* as described by Kemppainen et al. (2020). This OE construct incorporated the *Laccaria GPD* gene (JGIv2 ID #318873) Kozak sequence to improve transgene expression in *Laccaria* (Table **S1**, Fig **S1**). *Laccaria bicolor* S238N vegetative mycelium was transformed using *Agrobacterium*-mediated transformation (Kemppainen *et al.*, 2005). Twelve to fourteen transformed fungal lines per construct were selected with 300 μg/mL hygromycin B and thereafter were thereafter maintained under 200 μg/mL hygromycin B selection pressure on modified P5 medium.

**Methods S5** Characterization of T-DNA insertion number in transgenic *L. bicolor* lines by ddPCR

Alterations of target gene expression were determined by qRT-PCR in the free-living mycelia of 12-14 randomly selected independent *L. bicolor* transformant lines as compared to wild-type mycelium (Fig. **S2**). Of each construct, two or three fungal transformants with successfully altered gene expression were selected for analysing the T-DNA insertion number and the genomic integration site as well as for further molecular and physiological analyses. Genomic DNA was extracted from selected *L. bicolor* transgenic lines with DNeasy Plant Mini Kit (Cat. # 69104,

QIAGEN, Hilden, Germany) according to the manufacturer's instructions. About 60 ng of genomic DNA for each line was digested with HindIII (FD0505, Thermo Fisher Scientific, Waltham, Massachusetts, United States) in 25 µl reactions. T-DNA insertion number was characterized by digital droplet PCR (ddPCR) slightly modified from Głowacka *et al.* (2016) (Method **S7**). Genes amplified were the *Hygromycin b resistance gene* as target, and *Ubiquitin* and *Elongation Factor 3* as reference genes (Table **S1**).

**Method S6** Characterization of T-DNA insertion sites in transgenic *L. bicolor* by Plasmid rescue and TAIL-PCR

For the selected transgenic *L. bicolor* RNAi lines, plasmid rescue was performed according to Kemppainen *et al.* (2008), with the exception that BamHI (ER0051, Thermo Fisher Scientific, Waltham, Massachusetts, United States) was used to digest gDNA in the first step. TAIL-PCR and HiTAIL-PCR were performed for all overexpressor lines and for some RNAi strains, where plasmid rescue was not successful. The protocol and arbitrary degenerate (AD) primer were the same as described in Liu et al. (1995) and Liu et al. (2007). The sequences of left boundary primers L1, L2, L3 and long L2 are provided in Table **S1**. Bands were purified after gel-electrophoresis by QIAquick gel extraction kit (QIAGEN, Hilden, Germany) according to the manufacturer's instructions. Sequencing of plasmids from plasmid rescue and cut bands from TAIL-PCR and HiTAIL-PCR were performed with the Mix2Seq kit (Eurofins, Hvidovre, Denmark) using M13R primers (Table **S1**) for plasmid sequencing, and L2 or L3 primers for sequencing PCR products. The resulting sequences were identified in the *L. bicolor* genome using the BLASTN algorithm on the JGI genome portal (http://mycocosm.jgi.doe.gov/lacb2.home.html) in *L. bicolor* v2.0 masked assembly or *L. bicolor* v2.0 filtered model transcript databases. The T-DNA insertion number and the genomic insertion site annotations are provided in Table **S2**.

**Methods S7** Characterization of T-DNA insertion number in transgenic L. bicolor lines by ddPCR

The ddPCR reactions were prepared using 2 µL digested genomic DNA (about 5 ng), 0.22 µL forward and reverse primers (10 pmol/µL), 11 µL of commercial reaction mix (2x QX200 ddPCR EvaGreen Supermix, 186-4033, BioRad, Hercules, CA, USA), which includes an intercalating fluorescent dye, polymerase, $Mg^{2+}$ and dNTPs. The total reaction volume for each PCR was 22 µL. Droplets were generated with droplet generator cartridges, gaskets and cartridge holder (186-4007 and DG8, 186-3051, Bio-Rad,Hercules, CA,USA) in a droplet generator (QX200, 186-4002, Bio-Rad, Hercules, CA, USA). According to the manufacturer's instructions, 20 µL of ddPCR reaction mix and 70 µL of droplet generation oil (EvaGreen 186-4005, Bio-Rad, Hercules, CA, USA) were loaded into cartridges. The resulting 40 µL sample of generated droplets was dispensed in one well of a 96-well PCR semi-skirted plate (12001925, Bio-Rad, Hercules, CA, USA) and sealed with foil (Pierceable foil heat seal, 181-4040 and PX1 PCR plate sealer, 181-4000, Bio-Rad, Hercules, CA, USA). The droplet mix was cycled through a PCR program using a deep-well thermal cycler (T100, 186-1096, Bio-Rad, Hercules, CA, USA), followed immediately by analysis in a droplet reader (QX200, 186-4003, Bio-Rad, Hercules, CA, USA). Fluorescence reads per individual droplet from each well were analyzed with the manufacturer-provided software (Quanta Soft version 1.0, Bio-Rad, Hercules, CA, USA). For each reaction, two replicates were performed. The *HYGromycin B (HYGB)* resistance gene in the T-DNA was the target gene, the *LbUBiQuitin* (UBQ) gene and *LbELongation Factor 3 (ELF)* gene were the reference genes. Insertion numbers were calculated as ratios between *HYGB* and each reference gene. Primers are listed in Table **S1**.
